# Self-reporting transposons enable simultaneous readout of gene expression and transcription factor binding in single cells

**DOI:** 10.1101/538553

**Authors:** Arnav Moudgil, Michael N. Wilkinson, Xuhua Chen, June He, Alex J. Cammack, Michael J. Vasek, Tomas Lagunas, Zongtai Qi, Samantha A. Morris, Joseph D. Dougherty, Robi D. Mitra

## Abstract

*In situ* assays of transcription factor (TF) binding are confounded by cellular heterogeneity and represent averaged profiles in complex tissues. Single cell RNA-seq (scRNA-seq) is capable of resolving different cell types based on gene expression profiles, but no technology exists to directly link specific cell types to the binding pattern of TFs in those cell types. Here, we present self-reporting transposons (SRTs) and their use in single cell calling cards (scCC), a novel assay for simultaneously capturing gene expression profiles and mapping TF binding sites in single cells. First, we show how the genomic locations of SRTs can be recovered from mRNA. Next, we demonstrate that SRTs deposited by the *piggyBac* transposase can be used to map the genome-wide localization of the TFs SP1, through a direct fusion of the two proteins, and BRD4, through its native affinity for *piggyBac*. We then present the scCC method, which maps SRTs from scRNA-seq libraries, thus enabling concomitant identification of cell types and TF binding sites in those same cells. As a proof-of-concept, we show recovery of cell type-specific BRD4 and SP1 binding sites from cultured cells. Finally, we map Brd4 binding sites in the mouse cortex at single cell resolution, thus establishing a new technique for studying TF biology *in situ*.

## Introduction

Transcription factors (TFs) regulate gene expression during the most critical junctures in the specification of cell fate [1-4]. They are central to the maintenance of stem cell pluripotency [5,6] and required for normal organogenesis during development [7]. Overexpression of certain TFs can transdifferentiate one cell type into another [8], while abolishing TF binding sites can result in striking global phenotypes [9,10]. Furthermore, the pattern of TF binding is often dysregulated during disease states [11]. A better understanding of TF binding during tissue development and homeostasis would provide insights into how cellular diversity arises and is maintained under normal and abnormal biological conditions.

In the past few years, single cell RNA-seq (scRNA-seq) has emerged as the *de facto* approach for characterizing cellular diversity in complex tissues and organisms [12-17]. More recently, multi-modal scRNA-seq technologies have been developed [18-24] that combine transcriptional information with other genomic assays. These methods are motivated by the realization that while scRNA-seq can describe the current state of a biological system, it alone cannot explain how that state arose. Thus, for a given population of cells, one can now simultaneously measure transcriptome and genome [18,19], or methylome [20,21], or chromatin accessibility [21,22], or cell-surface markers [23,24]. These techniques enable greater insight into the regulatory elements driving individual transcriptional programs.

A notable lacuna in the single cell repertoire is a method for simultaneously assaying transcriptome and TF binding. Such a method would allow for the genome-wide identification of TF binding sites across multiple cell types in complex tissues. ChIP-seq is the most popular technique for studying TF binding [25], and while single cell ChIP-seq has been previously described [26], this technique has only been employed to map highly abundant proteins such as methylated histones. DamID can recover TF binding sites by identifying nearby exogenously methylated adenines [27,28], but in single cells it has only been used to study laminin-associated domains [29-31]. Importantly, both methods yield sparse data, either in small numbers of cells [31] or without simultaneously capturing mRNA [26]. Thus, each can only be used in heterogeneous samples if the cell type is known *a priori* and if sufficient numbers of cells are obtained by selection or sorting to overcome sparsity. In contrast, single cell assay for transposase-accessible chromatin (scATAC-seq) [32] can be used to identify nucleosome-free regions that may be bound by TFs across large numbers of mixed cells. However, it can only suggest potential DNA binding proteins by motif inference. Thus, it does not directly measure TF occupancy, and moreover it cannot be used to study transcriptional regulators that bind DNA indirectly or non-specifically, such as chromatin remodelers.

Our lab has previously developed transposon calling cards as an alternative assay of TF binding [33-35]. This system relies on two components: a fusion between a TF and a transposase, and a transposon carrying a reporter gene. The fusion transposase deposits transposons near TF binding sites; these insertions are subsequently amplified from genomic DNA and subjected to high-throughput sequencing. Thus, the redirected transposase leaves “calling cards” at the genomic locations it has visited, which can be identified later in time. The result is a genome-wide assay of all binding sites for that particular TF. In mammalian cells, we have heterologously expressed the *piggyBac* transposase [36] fused to the TF SP1 and shown that the resulting pattern of insertions reflects SP1’s DNA binding preferences [35]. However, the method was only feasible in bulk preparations.

Here we present single cell calling cards (scCC), an extension of transposon calling cards that simultaneously profiles mRNA abundance and TF binding at single cell resolution. The key component of our work is a novel construct called the self-reporting transposon (SRT). Using SRTs, the genomic locations of inserted transposons can be mapped from either mRNA or DNA, but the use of mRNA leads to both higher efficiency and compatibility with single-cell transcriptomics. We first establish that TF-directed SRTs, in bulk, retain the ability to accurately identify TF binding sites. Next, we demonstrate that the unfused *piggyBac* transposase, through its native affinity for the bromodomain TF BRD4, can be used to identify BRD4-bound super-enhancers (SEs). We then present the scCC method, which allows cell-specific mapping of SRTs from scRNA-seq libraries. This enables, in one experiment, concomitant assignment of cell types and identification of TF binding sites within those cells. As a proof-of-concept, we use scCC to map BRD4 and SP1 sites in mixtures of cultured human cells. We conclude by identifying cell type-specific Brd4 binding sites *in vivo* in the postnatal mouse cortex. These results demonstrate that scCC could be a broadly applicable tool for the study of specific TF binding interactions across all cell types within heterogeneous tissues.

## Results

### Self-reporting transposons can be mapped from mRNA instead of genomic DNA

In order to combine scRNA-seq with calling cards, we sought to develop a transposon whose genomic position could be determined from mRNA. We created a *piggyBac* self-reporting transposon (SRT) by removing the polyadenylation signal from our standard DNA-based calling card vector (Fig. 1A). This enables RNA polymerase II (Pol II) to transcribe the reporter gene contained in the transposon and continue through the terminal repeat (TR) into the flanking genomic sequence. Thus, SRTs “self-report” their locations through the unique genomic sequence found within the 3’ untranslated regions (UTRs) of the reporter gene transcripts. Although previously published gene-or enhancer-trap transposons could, in principle, also capture local positional information via RNA, they are resolution-limited to the nearest gene or enhancer, respectively [37]. In contrast, the 3’ UTRs of SRT-derived transcripts contain the transposon-genome junction in the mRNA sequence, so we can map insertions with base-pair precision.

**Figure 1:**
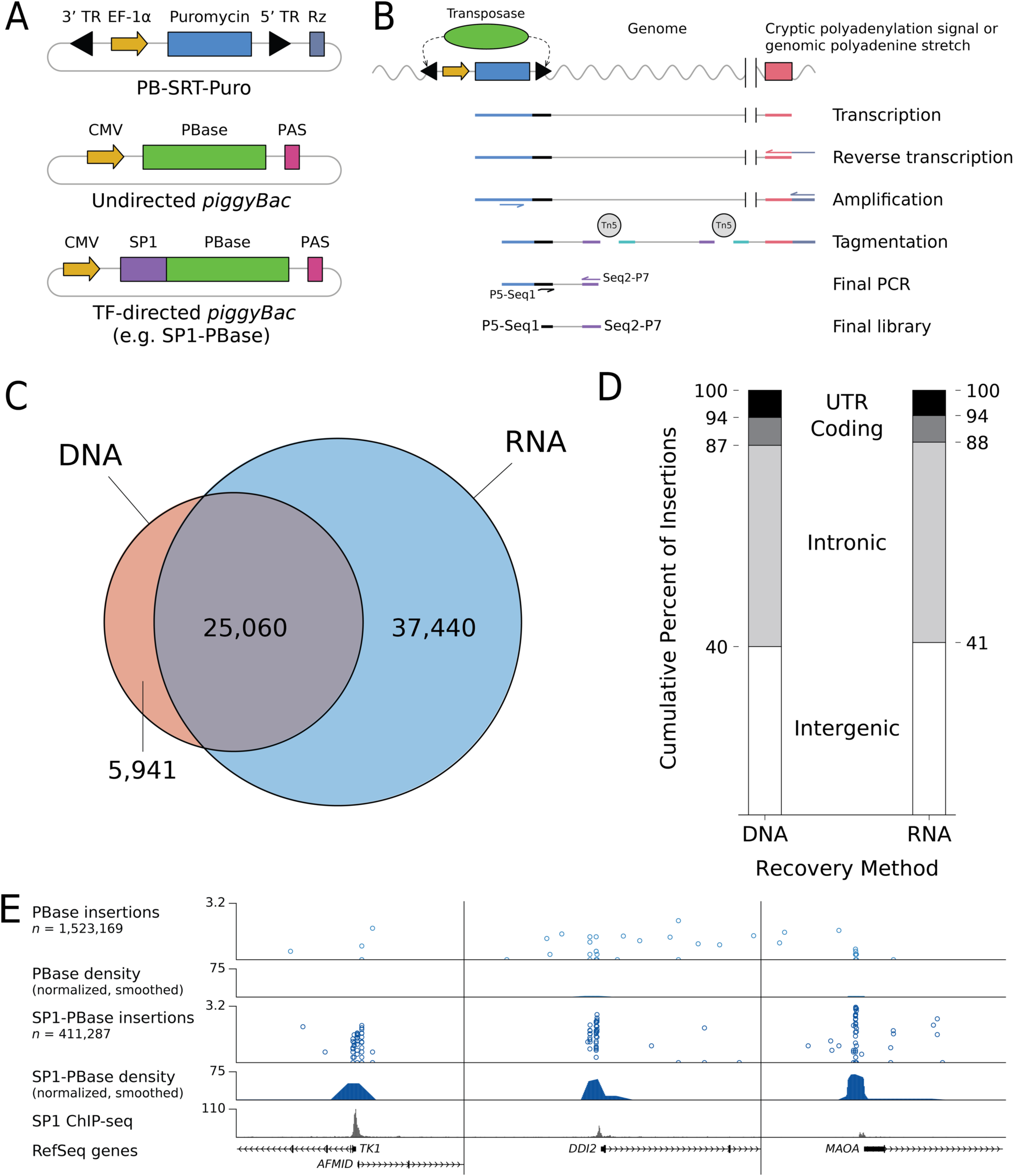
Self-reporting transposons (SRTs) are mapped more efficiently from RNA over DNA and, when directed SP1-PBase, identify SP1 binding sites. (A) Schematic of a self-reporting *piggyBac* transposon with puromycin marker (PB-SRT-Puro) and undirected (PBase) and SP1-directed (SP1-PBase) *piggyBac* transposases. SRTs are constructed by removing the PAS sequence between the end of the marker gene and the 5’ TR. A self-cleaving ribozyme (Rz) on the delivery vector, downstream of the SRT, prevents recovery of plasmid transposons. (B) SRTs are mapped by reverse transcribing RNA with a poly(T) primer followed by a series of nested PCRs and tagmentation. This final library is enriched for the junction between the transposon and the genome. (C) RNA-based recovery of SP1-directed SRTs in HCT-116 cells is more efficient than DNA-based recovery. The RNA protocol recovers 80% of the same insertions as the DNA protocol and recovers twice as many insertions overall. (D) The distribution of insertions with respect gene annotation is identical between transposons recovered by DNA and by RNA. (E) Insertions deposited by SP1-PBase show pronounced and specific clustering at SP1 ChIP-seq peaks over insertions left by undirected PBase. In the calling card track, each circle represents an independent insertion. Genomic position is on the *x*-axis and the number of reads supporting that insertion is on the *y*-axis on a *log_10_*-transformed scale. The density tracks show the local rate of insertions in each experiment (IPM/kb), normalized for library size, and smoothed in the WashU Epigenome Browser. TR: terminal repeat; Puro: puromycin; PAS: polyadenylation signal; IPM: insertions per million mapped insertions; kb: kilobase.

SRTs are mapped following reverse transcription (RT) and PCR amplification of self-reporting transcripts. These transcripts contain stretches of adenines that are derived from either cryptic polyadenylation signals (PAS) or polyadenine tracts encoded in genomic DNA downstream of the SRT insertion point (Fig. 1B). A poly(T) RT primer hybridizes with these transcripts and introduces a universal priming site at one end of the transcripts. A pair of nested PCRs with an intermediate tagmentation [38] step enable recovery of the transposon-genome junction. After adapter trimming and alignment, the 5’ coordinates of these reads identify the genomic locations of insertions in the library. Libraries generated without transposase produce very few genomically mapped reads but the protocol is highly efficient when transposase is added (Supp. Fig. 1A).

To compare transposon recovery between the new RNA-based protocol and our standard DNA-based inverse PCR protocol [35], we transfected HCT-116 cells with a plasmid carrying a *piggyBac* SRT (PB-SRT-Puro) and a plasmid encoding a fusion of the TF SP1 and *piggyBac* transposase (SP1-PBase; Fig. 1A). After two weeks of selection, we obtained approximately 2,300 puromycin-resistant clones. We split these cells in half: one half underwent inverse PCR while the other half were processed with our new RNA-based method. With inverse PCR, we obtained 31,001 insertions (mean coverage: 709 reads per insertion), while the RNA-based protocol recovered 62,500 insertions (mean coverage: 240 reads per insertion). About 80% of insertions recovered by DNA calling cards were also recovered in the RNA-based library (25,060 insertions; Fig. 1C), an overlap comparable to that between technical replicates of the RNA workflow (Supp. Fig. 1B). However, the RNA protocol recovered a further 37,440 insertions that were not found in the DNA-based library. To determine if these extra insertions were genuine, we analyzed the distribution of insertions by genetic annotation (Fig. 1D) or chromatin state (Supp. Fig. 1C; Supp. Table 1). Transposons mapped from either the DNA or the RNA libraries showed comparable distribution into annotated domains of particular functional or chromatin states, indicating that RNA recovery of transposons appears to be unbiased with respect to our established, DNA-based protocol.

Since *piggyBac* is known to preferentially insert near active chromatin [39], we wondered whether SRT recovery was biased towards euchromatic regions. Prior studies have shown that the *Sleeping Beauty* transposase [40,41] has very little preference for chromatin state [39]. We created a self-reporting *Sleeping Beauty* transposon and compared its genome-wide distribution to that of SRTs deposited by wild-type *piggyBac* (Supp. Fig. 2A-B). Undirected *piggyBac* transposases appeared to modestly prefer transposing into promoter and enhancers, which is consistent with previous reports [39,42] (Supp. Table 1). By contrast, *Sleeping Beauty* showed largely uniform rates of insertions across all chromatin states, including repressed and inactive chromatin (Supp. Fig. 2B). These results affirm that while RNA-based recovery is more efficient, it still preserves the underlying genomic distributions of insertions. Furthermore, because SRTs can be recovered from virtually any chromatin state, RNA-based calling card recovery can be employed to analyze a variety of TFs with broad chromatin-binding preferences.

A common artifact observed in DNA-based transposon recovery is a large fraction of reads mapping back to the donor transposon plasmid instead of the genome. Although this can be mitigated by long selection times or by digestion with the methyladenine-sensitive enzyme DpnI [35], these methods do not completely eliminate background and are not compatible with all experimental paradigms, viral transduction in particular. To reduce this artifact, we included a hammerhead ribozyme [43] in the SRT plasmid downstream of the 5’ TR. Before transposition, the ribozyme will cleave the nascent transcript originating from the marker gene, thus preventing RT. Transposition allows the SRT to escape the downstream ribozyme, leading to recovery of the self-reporting transcript. In our comparison of DNA-and RNA-based recovery, about 15% of reads from the SP1-PBase DNA library aligned to the plasmid, compared to fewer than 1% of reads from the RNA library (Supp. Fig. 1D). Thus, the addition of a self-cleaving ribozyme virtually eliminates recovery of un-excised transposons.

### SP1 fused to *piggyBac* directs SRT insertions to SP1 binding sites

We next sought to confirm that RNA calling cards, in bulk, can still be used to identify TF binding sites. We transfected 10-12 replicates of HCT-116 cells with plasmids containing the PB-SRT-Puro donor transposon and SP1 fused to either *piggyBac* (SP1-PBase) or a hyperactive variant of *piggyBac* [44] (SP1-HyPBase). As controls, we also transfected a similar number of replicates with undirected PBase or HyPBase, respectively. We obtained 411,287 insertions from SP1-PBase and 1,523,169 insertions from PBase. Similarly, we obtained 2,033,229 SP1-HyPBase insertions and 5,779,101 insertions from HyPBase.

Fig. 1E and Supp. Fig. 4A show the redirection of SRT calling cards by SP1-PBase and SP1-HyPBase, respectively, to three representative SP1-bound regions of the genome. Each circle in the insertions track is an independent transposition event whose genomic position is on the *x*-axis. The *y*-axis is the number of reads supporting each insertion on a *log_10_* scale. To better compare transposition across libraries with different numbers of insertions, we plotted the normalized local insertion rate as a density track. All three of the loci depicted in Fig. 1E and Supp. Fig. 4A show a specific enrichment of calling card insertions in the SP1 fusion experiments that is not observed in the undirected control libraries. Next, we called peaks at all genomic regions enriched for SP1-directed transposition. The number of insertions observed at significant peaks for both SP1-PBase and SP1-HyPBase was highly reproducible between biological replicates (*R^2^* = 0.84 and 0.96, respectively; Supp. Fig. 3A and Supp. Fig. 4B). Furthermore, calling card peaks were highly enriched for SP1 ChIP-seq signal at their centers, both on average (Supp. Fig. 3B and Supp. Fig. 4C) and in aggregate (Supp. Fig. 3C and Supp. Fig. 4D). SP1 is known to preferentially bind near TSSs [45,46] and is also thought to play a role in demethylating CpG islands [47-49]. We confirmed that the SP1-directed transposases preferentially inserted SRT calling cards near TSSs, CpG islands, and unmethylated CpGs at statistically significant frequencies (*p* < 10^−9^ in each instance, *G* test of independence; Supp. Fig. 3D and Supp. Fig. 4E). Moreover, compared to undirected *piggyBac*, SP1-directed *piggyBac* showed a striking preference for depositing insertions into promoters (Supp. Fig. 2A-B). Lastly, regions targeted by SP1-PBase and SP1-HyPBase were enriched for the canonical SP1 DNA binding motif (*p* < 10^−70^ in each instance; Supp. Fig. 3E and Supp. Fig. 4F). Taken together, these results indicate that SP1 can redirect *piggyBac* SRTs near SP1 binding sites.

### Clustering of undirected *piggyBac* insertions identifies BRD4-bound super-enhancers

Previous studies have shown that the undirected *piggyBac* transposase preferentially inserts transposons near super-enhancers (SEs) [39], a unique regulatory element that is thought to play a critical role in regulating cell identity [50]. SEs are often enriched for the histone modification H3K27ac as well as RNA polymerase II and general transcription factors like the mediator element MED1 and the bromodomain protein BRD4 [50-52]. Moreover, the *piggyBac* transposase has a strong biophysical affinity for BRD4, as these proteins can be co-immunoprecipitated [42]. We hypothesized that, given the millions of insertions we assayed from the undirected PBase and HyPBase controls in the SP1-directed experiments (Fig. 1E, Supp. Fig. 4A), we would be able to identify BRD4-bound SEs simply from the localization of undirected *piggyBac* transpositions.

Both undirected PBase and HyPBase showed non-uniform densities of insertions at loci bound by BRD4 (Fig. 2A, Supp. Fig. 7). At statistically significant peaks of *piggyBac* calling cards, PBase and HyPBase showed high reproducibility of normalized insertions between biological replicates (Fig. 2B, Supp. Fig. 5B). Next, we calculated the mean BRD4 enrichment, as assayed by ChIP-seq [53], across these peaks. *piggyBac* peaks showed significantly increased BRD4 signal compared to a genome-wide permutation of the peaks (*p* < 10^−9^ in both instances, Kolmogorov-Smirnov test; Fig. 2C and Supp. Fig. 5C). Maximum BRD4 ChIP-seq signal was observed at calling card peak centers and decreased symmetrically in both directions. We also found that *piggyBac* peaks show striking ChIP-seq patterns for several histone modifications [54,55], in particular an enrichment for H3K27ac ChIP-seq signal (Fig. 2D, Supp. Fig. 5D). Since bromodomains bind acetylated histones, this observation further supports the hypothesis that undirected *piggyBac* insertions can be used to map BRD4 binding. These peaks were also enriched in H3K4me1, another canonical enhancer mark, and depleted for H3K9me3 and H3K27me3, modifications associated with repressed chromatin [56]. Taken together, these results demonstrate that *piggyBac* insertion density is highly correlated with BRD4 binding throughout the genome and that regions enriched for undirected *piggyBac* insertions share features common to enhancers.

**Figure 2:**
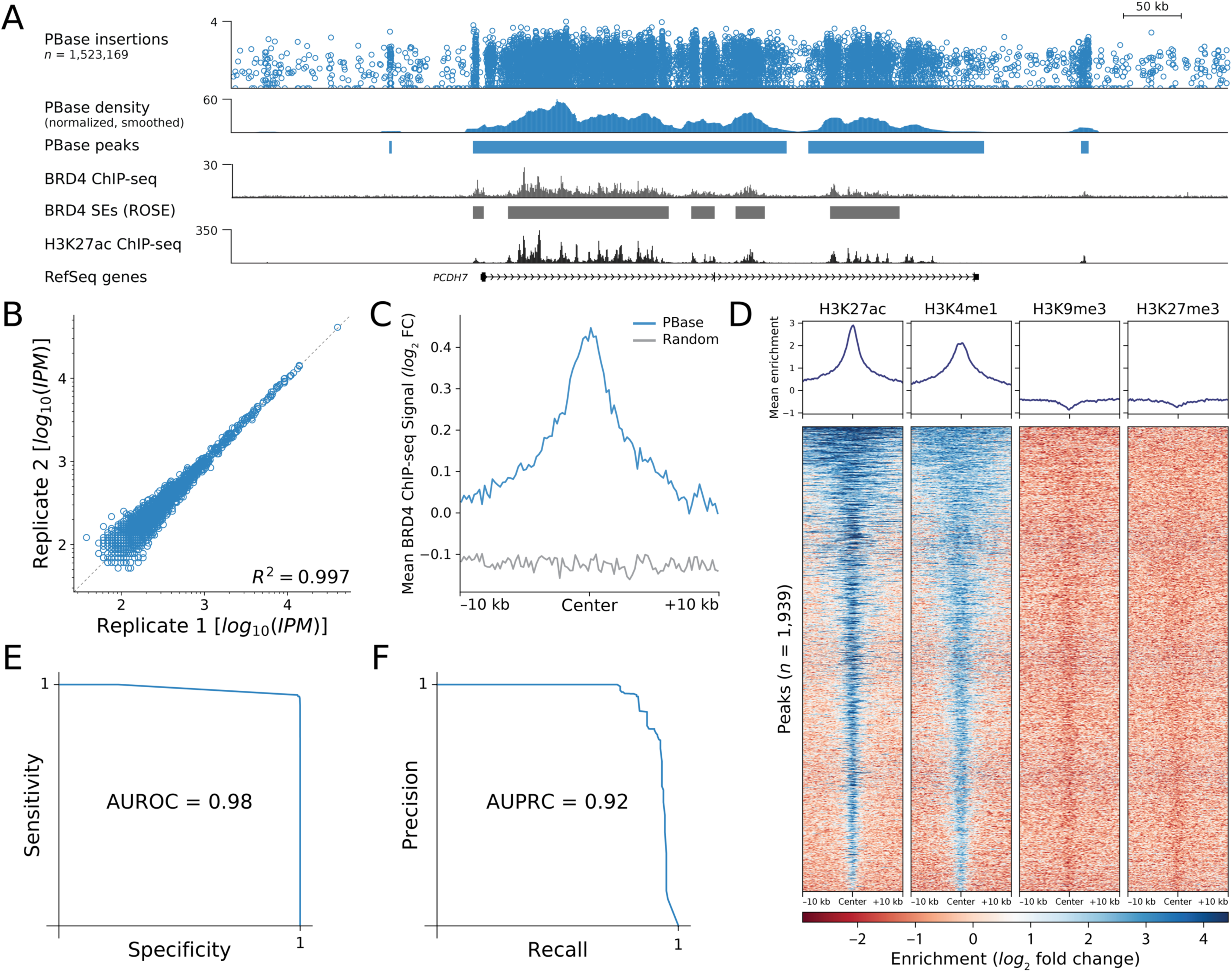
Undirected *piggyBac* (PBase) insertions mark BRD4-bound super-enhancers. (A) Undirected PBase insertions are distributed non-randomly, with increased density overlapping BRD4-bound chromatin and H3K27 acetylated histones. Also shown are BRD4-bound SEs. (B) PBase peak calls are highly replicable, with biological replicates showing high concordance of normalized insertions at peaks. (C) PBase peaks show central enrichment for BRD4 ChIP-seq signal. These findings are statistically significant when compared to a genome-wide permutation of PBase peaks (*p* < 10^−9^, KS test). (D) PBase peaks are centrally enriched for the histone modifications H3K27ac and H3K4me1, marks associated with enhancers. These same peaks show mild depletion for H3K9me and H3K27me, marks canonically associated with repressed chromatin. (E) Receiver-operator characteristic curve for SE detection using PBase peaks. (F) Precision-recall curve for SE detection using PBase peaks. SE: super-enhancer; IPM: insertions per million mapped insertions; AUROC: area under receiver-operator curve; AUPRC: area under precision-recall curve; KS: Kolmogorov-Smirnov; FC: fold change.

We next assessed whether *piggyBac* peaks can be used to identify BRD4-bound SEs. We used BRD4 ChIP-seq data from HCT-116 cells [53] to create a reference list of BRD-bound SEs [51,57] (Fig. 2A, Supp. Fig. 5A). We then constructed receiver-operator characteristic curves based on our ability to detect SEs from PBase-and HyPBase-derived peaks (Fig. 2E and Supp. Fig. 5E). The high areas under the curve (0.98 in each instance) indicate that we can robustly identify BRD4-bound SEs from *piggyBac* transpositions. Across a range of sensitivities, calling card peaks are highly specific and have high positive predictive value (AUPRC = 0.92 in each instance; Fig. 2F and Supp. Fig. 5F). Thus, undirected *piggyBac* transpositions are an accurate assay of BRD4-bound SEs.

To better understand the relationship between SE sensitivity and the number of insertions recovered, we downsampled the data from the PBase and HyPBase experiments in half-log increments (Supp. Fig. 6A-B). These heatmaps show that sensitivity increases with the total number of insertions recovered. Since we cannot predict how many, or few, insertions future experiments will yield, we also performed linear interpolation on the downsampled data. The resulting contour plots (Supp. Fig. 6C-D) indicate the approximate sensitivity of BRD4-bound SE detection in HCT-116 cells. Our analysis suggests that even with as few as 10,000 insertions, we can still obtain sensitivities around 50%.

### Single cell calling cards enables simultaneous identification of cell type and cell type-specific TF binding sites

We next sought to recover SRTs from scRNA-seq libraries. This would enable us to identify cell types from transcriptomic clustering and, using the same source material, profile TF binding in those cell types. We adopted the 10x Chromium platform given its high efficiency of cell and transcript capture as well as its ease of use [58]. Like many microfluidic scRNA-seq approaches [59,60], the cell barcode and unique molecular index (UMI) are attached to the 3’ ends of transcripts. This poses a molecular challenge for SRTs since the junction between the transposon and the genome may be many kilobases away, precluding the use of high-throughput short read sequencing. To overcome this barrier, we developed a circularization strategy to physically bring the cell barcode in apposition to the insertion site (Fig. 3A).

**Figure 3:**
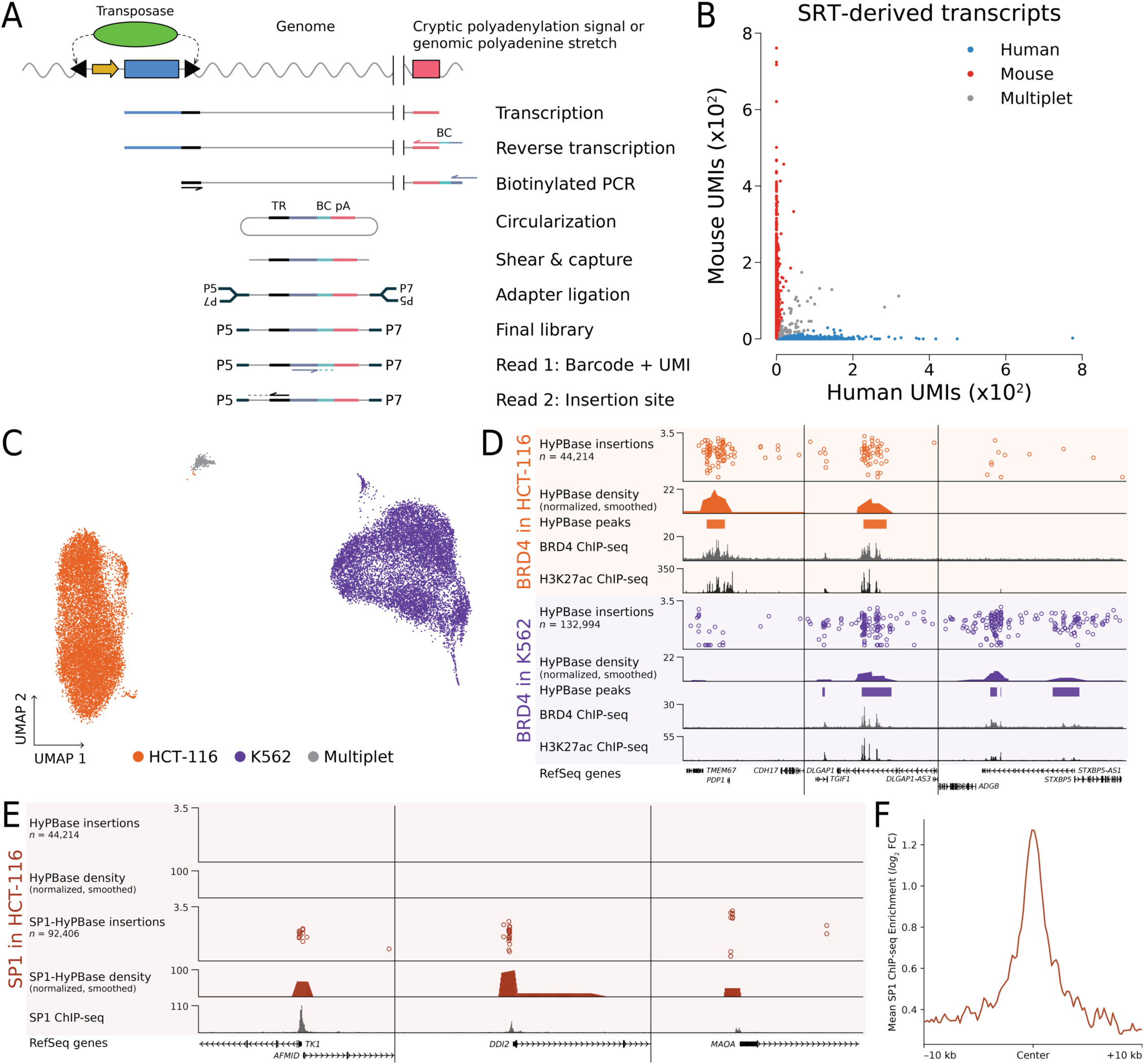
Single cell calling cards (scCC) maps BRD4 binding and SP1 in single cells. (A) Schematic of the scCC library preparation strategy from scRNA-seq libraries. Self-reporting transcripts are amplified using biotinylated primers and circularized, which brings the cell barcode and UMI in close proximity to the transposon-genome junction. Circularized molecules are sheared, captured with streptavidin, and Illumina adapters are ligated. Custom sequencing yields the cell barcode and UMI with read 1 and the genomic insertion site with read 2. (B) Barnyard plot of HCT-116 and N2a cells transfected with SRTs shows clean segregation of cell types. Most cells were assigned either human insertions or mouse insertions, with an estimated multiplet rate of 7.8%. (C) Human HCT-116 and K562 cells were transfected with PB-SRT-Puro and HyPBase and subsequently subjected to scRNA-seq. Two clear cell types emerge revealing each constituent cell population. (D) scCC deconvolves HyPBase insertions from HCT-116 and K562 cells, identifying shared and specific BRD4 binding sites. (E) scCC on HCT-116 cells transfected with SP1-HyPBase identifies SP1 binding sites. (F) SP1-HyPBase peaks from scCC data show strong central enrichment for SP1 ChIP-seq signal. TR: terminal repeat; BC: barcode; pA: poly(A) sequence; UMI: unique molecular index; Puro: puromycin; FC: fold change.

We used a modified version of the bulk SRT amplification protocol where we amplified with primers that bound to the universal priming sequence next to the cell barcode and the terminal sequence of the *piggyBac* TR. These primers were biotinylated and carried a 5’ phosphate group. The PCR products of this amplification were diluted and allowed to self-ligate overnight. They were then sheared and captured with streptavidin-coated magnetic beads. The rest of library was prepared on-bead and involved end repair, A-tailing, and adapter ligation. A final PCR step added the required Illumina sequences for high-throughput sequencing. The standard Illumina read 1 primer sequenced the cell barcode and UMI, while a custom read 2 primer, annealing to the end of the *piggyBac* 5’ TR, sequenced into the genome. Thus, we collected both the location of a *piggyBac* insertion as well as its cell of origin. We call this method single cell calling cards (scCC).

We validated the method by performing a species-mixing experiment using human HCT-116 cells and mouse N2a cells. Cells were mixed prior to droplet generation and the resulting emulsion was processed through first strand synthesis. At this point, half of the RT product was amplified according to the standard 10x protocol. The resulting scRNA-seq revealed strong species separation with an estimated multiplet rate of 3.2% (Supp. Fig. 8A). The remainder of the first strand synthesis was used for the scCC protocol. We restricted our calling card analysis to those insertions whose cell barcodes were observed in the scRNA-seq library. The distribution of insertions across these cells reflected a continuum from pure mouse to pure human (Supp. Fig. 8B-C). Since intramolecular ligation and subsequent PCR may introduce unwanted artifacts, such as mis-assignment of a barcode from cell type A to an insertion site in cell type B, we required that a given insertion in a given cell must have at least two different UMIs associated with it. Imposing this filter improved the number of pure mouse and human cells (Supp. Fig. 8D), yielding clear species separation with an estimated multiplet rate of 7.8% (Fig. 3B). This establishes that our method can accurately map calling card insertions in single cells.

We then asked whether scCC could discern cell type-specific TF binding. We transfected two human cell lines, HCT-116 and K562, with HyPBase and PB-SRT-Puro and mixed them together. The resulting scRNA-seq libraries clearly identified the two major cell populations (Fig. 3C; Supp. Fig. 9A). We then prepared scCC libraries from these cells and used the cell barcodes from the HCT-116 and K562 clusters to assign insertions to the two different cell types. We obtained 44,214 insertions from 12,891 HCT-116 cells (mean 3.4 insertions per cell; mean 136 reads per insertion) and 132,994 insertions from 11,912 K562 cells (mean 11 insertions per cell; mean 103 reads per insertion). The distribution of insertions per cell varied by cell type (Supp. Fig. 9D) and does not appear to be correlated with differences in total RNA content (Supp. Fig. 9B-C). Over 93% and 97% of HCT-116 and K562 cells, respectively, had at least one insertion event. Using scCC insertion data alone, we called peaks and successfully identified BRD4-bound loci that were specific to HCT-116 cells, shared between HCT-116 and K562, and specific to K562 cells, respectively (Fig. 3D). Both HCT-116 and K562 peaks showed statistically significant enrichment for BRD4 ChIP-seq signal (*p* < 10^−9^ in both instances, Kolmogorov-Smirnov test; Supp. Fig. 9E-F). From our earlier downsampling analysis, we estimated that with a *p*-value cutoff of 10^−9^, our sensitivity for detecting BRD4-bound SEs would be approximately 60% (Supp. Fig. 6D). The actual sensitivity at this level of recovery was 64%, validating that downsampling analysis can reasonably estimate the performance of scCC. In all, these experiments demonstrate that scCC can be used to deconvolve cell type-specific TF binding.

Since these Brd4 binding sites were identified using undirected HyPBase, we also sought to confirm that TF-*piggyBac* fusions would still work with scCC. We transfected HCT-116 cells with SP1-HyPBase and then performed scRNA-seq. We made scCC libraries from these experiments and identified 92,406 insertions from 30,682 cells (mean 3 insertions per cell; mean 129 reads per insertion). Over 84% of cells had at least one insertion. While previous studies have reported decreased activity of TF-*piggyBac* fusions [61], we observed similar distributions of insertions recovered per cell between HyPBase and SP1-HyPBase (Supp. Fig. 9G). As was observed in bulk (Supp. Fig. 4A), SP1-HyPBase-directed insertions recovered from single cells localize to SP1 binding sites (Fig. 3E). Finally, we investigated the reproducibility of the scCC method. Both single cell HyPBase and SP1-HyPBase showed high concordance between biological replicates at statistically significant peaks (Supp. Fig. 9H-I). Collectively, these experiments establish that scCC can be used to identify cell type-specific binding sites of both bromodomain and DNA-binding TFs.

**Figure 4:**
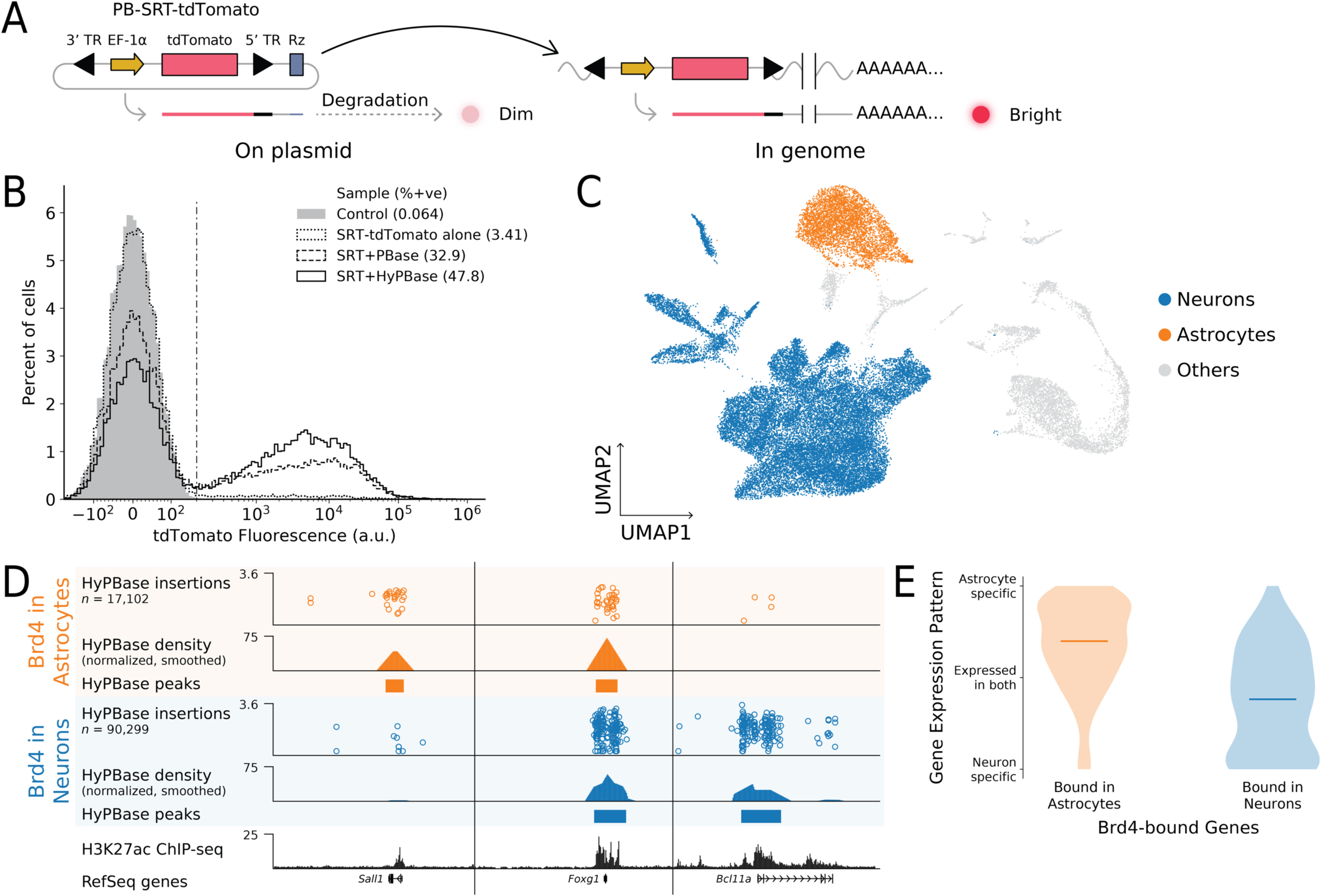
Single cell calling cards deconvolves Brd4-bound loci in the mouse cortex. (A) Schematic of PB-SRT-tdTomato, an SRT compatible with *in vivo* experiments. The pre-transposition tdTomato transcript (left) is degraded by the downstream ribozyme (Rz), leading to low fluorescence intensity. After transposition into the genome, the self-reporting transcript is stabilized and results in a bright signal. (B) Validation of PB-SRT-tdTomato in K562 cells. Cells transfected with both SRT and transposase (either PBase or HyPBase) show bimodal fluorescence enabling sorting for cells with insertions. (C) scRNA-seq analysis of mouse cortex libraries transduced with PB-SRT-tdTomato and HyPBase reveals multiple cell types, including astrocytes (*n* = 4,727) and neurons (*n* = 25,158). (D) scCC analysis of HyPBase insertions in astrocytes and neurons identify shared and specific Brd4 binding sites. Whole cortex H3K27ac ChIP-seq shown for comparison. (E) Gene expression specificity of genes overlapping astrocyte or sensitivity-matched neuron peaks. Expression values were taken from bulk RNA-seq. Specificity for each gene was calculated by dividing the expression of the gene in astrocytes by the sum of the expression values in astrocytes and neurons. Horizontal lines indicate the medians of the distributions. TR: terminal repeat.

### Single cell calling cards deconvolves cell type-specific Brd4 binding sites in the mouse cortex

To establish broad utility for scCC, we sought to record TF binding *in vivo*. Since *in vivo* models preclude puromycin selection, we designed an SRT carrying the fluorescent reporter tdTomato (Fig. 4A) and tested this reagent in cell culture. When this construct was transfected without transposase, 3.4% of cells register as tdTomato-positive, likely due to the action of the self-cleaving ribozyme downstream of the transposon. However, when the construct was co-transfected with PBase or HyPBase, this figure rose to 33% and 48%, respectively, corresponding to 11-and 16-fold increases in signal (Fig. 4B). In addition, cells transfected with only the fluorescent SRT produced very few reads that mapped to the genome, while the overwhelming majority of reads from cells co-transfected with transposase mapped to genomic insertions (Supp. Fig. 1A). Thus, this new construct, PB-SRT-tdTomato, allows us to select cells carrying calling card insertions by fluorescence activated cell sorting (FACS).

We chose the mouse cortex for our *in vivo* proof-of-concept because it is a heterogeneous tissue that has been the focus of several recent single cell studies [12,62-65]. We separately packaged the PB-SRT-tdTomato and HyPBase constructs in AAV9 viral particles [66] and delivered mixtures of both viruses to the developing mouse cortex via intracranial injections at P1. After 2-4 weeks, we dissected the cortex, dissociated it to a single cell suspension, performed FACS to isolate tdTomato-positive cells, and analyzed these cells by scRNA-seq and scCC using the 10x Chromium platform. We collected nine libraries in total comprising 35,950 cells and 113,859 insertions (Supp. Table 2). We clustered cells by their mRNA profiles and used established marker genes to classify different cell types (Supp. Fig. 10A-B) [63-65]. The two major cell populations recovered were neurons and astrocytes (Fig. 4C, Supp. Table 2), which is consistent with the known tropism of AAV9 [67]. We also identified a spectrum of differentiating oligodendrocytes and trace amounts of microglial, vascular, and ependymal cells. We then used the cell barcodes shared between the scRNA-seq and scCC libraries to assign insertions to specific cell types.

To determine whether scCC could recover biological differences between cell types *in vivo*, we analyzed HyPBase insertions in neurons and astrocytes, excluding neuroblasts and astrocyte-neuron doublets. We collected 90,299 insertions from 25,158 neurons and 17,102 insertions across 4,727 astrocytes. We then called peaks on the insertions within each cluster and identified astrocyte-specific, neuron-specific, and shared Brd4 binding sites (Fig. 4D). Brd4 ChIP-seq has not been reported for the mouse brain, but as Brd4 is known to bind acetylated histones, we compared our peak calls to a recent cortical H3K27ac ChIP-seq dataset [68]. Although this dataset was agglomerated over all cell types in the brain, we nevertheless found that peaks in both astrocytes and neurons showed statistically significant enrichment of H3K27ac signal (Supp. Fig. 11A, C; Kolmogorov-Smirnov *p*-value < 10^−9^ in each case). Brd4 is also thought to mark cell type-specific genes, so we identified genes that overlapped or were near astrocyte or neuron peaks and evaluated the specificity of expression of these genes. We identified 399 genes near astrocyte peaks and 211 genes near neuron peaks. We used bulk RNA-seq data from purified populations of cells [69] to assign gene expression values for each gene and plotted the distribution of these values along a continuum from purely astrocytic expression to purely neuronal expression. Genes near astrocyte peaks were more likely to be specifically expressed in astrocytes, and vice-versa for genes near neuron peaks (Fig. 4E). Gene Ontology enrichment analysis [70] on the astrocyte gene list included “gliogenesis,” and “glial cell differentiation,” as well as copper metabolism (Supp. Fig. 11B), a known function of astrocytes [71]; while the neuronal gene list was enriched for terms related to synapse assembly and neuron development (Supp. Fig. 11D). Overall, we conclude that scCC can accurately identify cell type-specific Brd4 binding sites *in vivo*.

Finally, we wondered if scCC could discriminate Brd4 binding between closely related cell types. From our scRNA-seq data (Fig. 5B; Supp. Fig. 10A-B), we identified upper and lower layer cortical excitatory neurons and compared HyPBase scCC data between them to identify shared and specific Brd4-bound loci (Fig. 5A). From 9,083 upper cortical neurons we obtained 30,225 insertions, which was on par with the 32,434 insertions collected from 6,980 lower cortical neurons. As a positive control, we identified a shared Brd4 binding site at the *Pou3f3* (*Brn-1*) locus (Fig. 5A, *p* < 10^−9^). *Pou3f3* was broadly expressed in both populations (Fig. 5C) and has been used to label layers 2-5 of the postnatal cortex [72,73]. We then identified differentially-bound regions in each cluster using insertions from the other cluster as a control. Upper cortical neurons showed specific Brd4 binding at *Pou3f2* (*Brn-2*), which is more restricted to layers 2-4 than *Pou3f3* [73,74], while lower cortical neurons showed Brd4 binding at *Bcl11b* (*Ctip2*) and *Foxp2*, common markers of layer 5 and layer 6 neurons, respectively (Fig. 5A; *p* < 10^−9^ in each instance) [73,75]. The expression patterns of these genes mirrored Brd4’s binding specificity, with *Pou3f2’*s expression mostly retained to the layer 2-4 cluster and the expression of *Bcl11b* and *Foxp2* restricted to the layer 5-6 neuron population (Fig. 5C). This demonstrates that scCC can identify differentially bound loci between very similar cell types.

**Figure 5:**
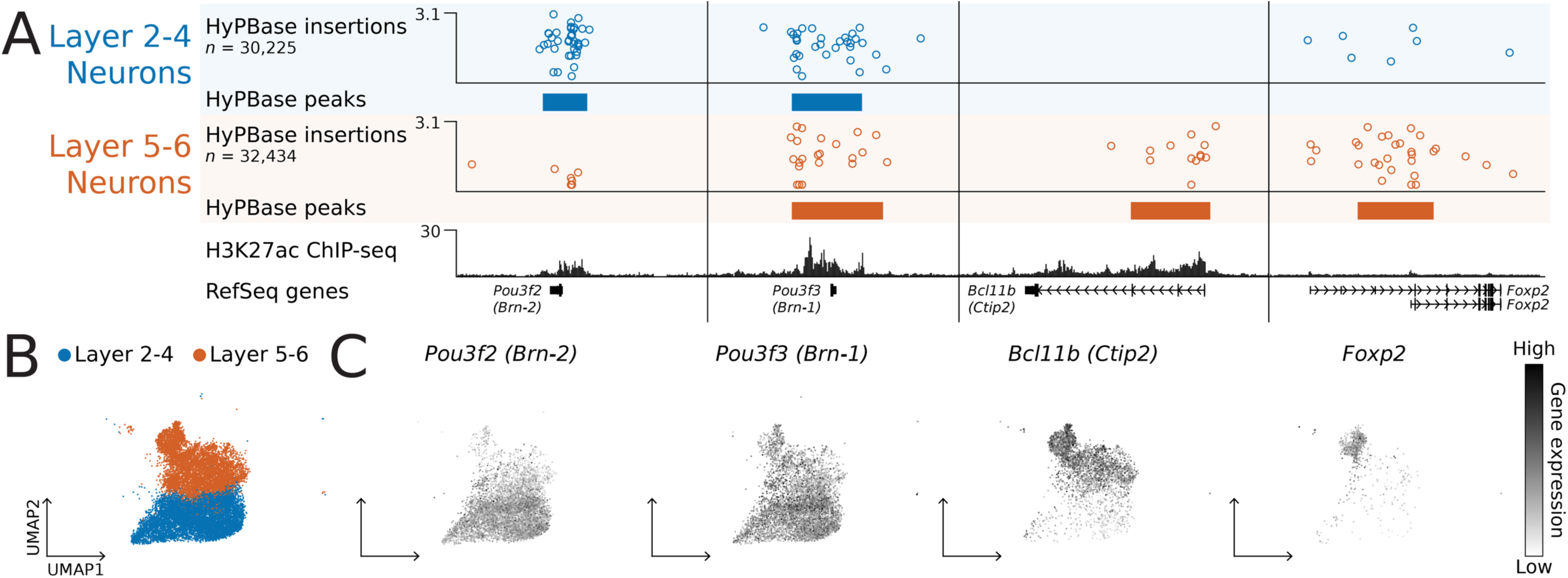
Single cell calling cards deconvolves Brd4 binding in cortical excitatory neurons and identifies known layer markers. (A) scCC analysis of HyPBase insertions in upper (layer 2-4) or lower (layer 5-6) cortical excitatory neurons identifies shared and specific Brd4 binding sites. Whole cortex H3K27ac ChIP-seq shown for comparison. (B) Layer 2-4 (*n* = 9,083) and layer 5-6 (*n* = 6,980) cortical excitatory neurons highlighted among the scRNA-seq clusters. (C) Gene expression patterns of the four genes from (A) mirrors the cell type-specificity of Brd4 binding.

## Discussion

Mapping TF binding in heterogeneous tissues is a challenging problem because traditional methods combine signals from multiple cell types into a single, agglomerated profile. The difficulty is further compounded if individual cell types are difficult to identify, isolate, or are rare, precluding their study. Single cell RNA-seq is a promising paradigm for handling such heterogeneity. Until now, it has been impossible to directly study the actions of individual TFs and connect them to specific cell states. We have presented a new method, single cell calling cards (scCC), that enables simultaneous identification of cell types and TF binding sites from complex mixtures and tissues. This is an important addition to the single cell repertoire and fills a recognized void in the field [76,77]. We anticipate this technique will enable researchers to study the consequences of TF binding in a variety of *ex vivo* and *in situ* models.

A concern with any transposon-based technique is the potential for deleterious interruption of target genes leading to cell death and, consequently, false negatives. Previous experiments in diploid yeast found that calling cards are deposited into promoters of essential and non-essential genes at comparable frequencies [34]. Since mammalian genomes have much larger intergenic regions than yeast, human and mice genomes are likely also able to tolerate calling card transpositions. Indeed, that we were able to deposit SRTs in the developing mouse brain into enhancers and super-enhancers suggests a small mutagenic burden.

One of the limitations of this technique is the relatively few insertions recovered on a per-cell basis, inflating the number of cells that must be analyzed to achieve good sensitivity. Previous studies have reported up to 15-30 insertions per cell for PBase [78-81], and likely higher for HyPBase [44,82]. While we observed similar performance from bulk RNA, we recovered fewer insertions per cell than this, on average, in our single cell experiments. This is likely due to the low capture rate of mRNA transcripts, which is common to all scRNA-seq methods [83]. The inclusion of cis-regulatory features known to enhance mRNA maturation and stability, such as the woodchuck hepatitis virus post-transcriptional regulatory element (WPRE) may increase representation of SRTs in scRNA-seq libraries. Furthermore, as the transcript capture rates of scRNA-seq technologies improve, we expect the sensitivity of our method will increase. The sensitivity of scCC can also be improved by simply analyzing larger numbers of cells, such as with Cell Hashing [84] or combinatorial barcoding [62]. Since the per-cell costs for scRNA-seq are exponentially falling [85], we expect that scCC can be used to analyze TF binding in even very rare cell types in the near future.

Our scCC experiments employed the *piggyBac* transposase, but for some applications, other transposases may prove advantageous. *piggyBac* inserts almost exclusively into TTAA tetranucleotides. For TFs that bind GC-rich regions or have high GC-content motifs, *piggyBac* fusions may have a difficult time finding nearby insertion sites. *Sleeping Beauty*, which inserts into TA dinucleotides, or *Tol2*, which does not have a strict insertion site preference [39], could be used to overcome these limitations. However, the natural affinity of the *piggyBac* transposase for BRD4 makes it the ideal choice for the study of BRD4-bound SEs, which play important regulatory roles in development and disease [52]. It is unclear why *piggyBac* shows such an affinity. BRD4 has an intrinsically disordered region and cooperative interactions between BRD4 and coactivators like MED1 may mediate the formation intranuclear condensates [86] at SEs. One hypothesis is that *piggyBac* has a similarly disordered domain that allows it to preferentially enter condensates and enrich SEs with insertions.

The defining feature of the scCC method is the self-reporting transposon (SRT). While here we have reported the *piggyBac* and *Sleeping Beauty* SRTs, the self-reporting paradigm should be generalizable to any transposon lacking a polyadenylation signal (PAS) in at least one terminal repeat. Expanding the palette of SRTs will illuminate the genome-wide behaviors of transposases and may yield further insight into chromatin dynamics [39]. Simultaneous expression of many TFs, each tagged to a different transposase, may also enable multiplexed studies of TF binding in the same cells. Mapping SRTs using cellular RNA appears to be substantially more efficient than the DNA-based inverse PCR method, but the reasons for this are unclear. Some efficiency is likely gained by eliminating self-ligation, as well as having multiple mRNA copies of each insertion to buffer against PCR artifacts. It is also unknown what fraction of self-reporting transcripts are actually polyadenylated as opposed to merely containing A-rich genomic tracts. Non-genic PASs prevent anti-sense transcription [87], which suggests that PASs may be more common in the genome than previously appreciated. Targeted 3’-end sequencing [88,89] of SRT libraries should help resolve this question, while long-read sequencing of self-reporting transcripts may identify non-canonical PASs. Finally, SRTs could lead to new single cell transposon-based assays. For example, just as CRISPR/Cas9 has been combined with scRNA-seq to read out the transcriptional effects of gene deletion [90,91], SRTs will allow transposon mutagenesis screens to be read out by scRNA-seq in a highly parallel fashion.

Finally, as calling card insertions are genomically integrated and preserved through mitosis, they could serve as records of molecular memory. The use of an inducible transposase [92] would enable the recording and identification of temporally-restricted TF binding sites. This would help uncover the stepwise order of events underlying the regulation of specific genes and inform cell fate decision making. More generally, transposon insertions could serve as barcodes of developmental lineage. Single transposition events have been used to delineate relationships during hematopoiesis [93,94]. Multiplexing several SRTs across every cell in an organism could code lineage in a cumulative and combinatorially diverse fashion, generating high-resolution cellular phylogenies.

## Acknowledgements

We would like to thank Jessica Hoisington-Lopez and MariaLynn Crosby from the DNA Sequencing Innovation Lab at The Edison Family Center for Genome Sciences and Systems Biology for their sequencing expertise. Additional sequencing services were performed by the Genome Technology Access Center in the Department of Genetics at Washington University School of Medicine. The Center is partially supported by NCI Cancer Center Support Grant P30 CA91842 to the Siteman Cancer Center and by ICTS/CTSA Grant UL1 TR000448 from the National Center for Research Resources (NCRR), a component of the National Institutes of Health (NIH), and NIH Roadmap for Medical Research. This work was also supported by the Hope Center Viral Vectors Core and a P30 Neuroscience Blueprint Interdisciplinary Center Core award to Washington University (P30 NS057105). Finally, we would like to thank Donald Conrad and Ben Humphreys for their advice and constructive feedback during this project.

This publication is solely the responsibility of the authors and does not necessarily represent the official view of NCRR or NIH.

## Author Contributions

A.M., M.N.W., Z.Q., and R.D.M. developed the self-reporting transposon (SRT) technology. A.M. and M.N.W. created and optimized the molecular workflow for recovering SRTs from bulk RNA-seq libraries, with contributions from Z.Q. A.M. developed and optimized the protocol for recovering SRTs from single cell RNA-seq libraries. S.A.M. provided guidance and assistance with single cell experiments. Z.Q. and T.L. cloned SRT constructs. A.M., M.N.W., J.D.D., and R.D.M. designed the experiments. A.M., X.C., and J.H. performed the *in vitro* experiments. M.N.W., A.J.C., and M.J.V performed the *in vivo* experiments. A.M., X.C., and J.H. generated sequencing libraries. A.M., J.D.D., and R.D.M. analyzed the data. A.M. and R.D.M. wrote the manuscript in consultation with all authors.

## Methods

### Cell culture

HCT-116, N2a, and HEK293T cells were cultured in Dulbecco’s Modified Eagle Medium (DMEM; Gibco #11965-084) supplemented with 10% fetal bovine serum (FBS; Peak Serum #PS-FB3) and 1% antibiotic-antimycotic (Anti-Anti; Gibco #15240-062). K562 cells were grown under the same conditions as the HCT-116 and N2a except replacing DMEM with RPMI 1640 Medium (Gibco #11875-085). Cells were grown at 37°C with 5% carbon dioxide (CO2). Puromycin (Sigma #P8899) was added 24 hours after transfection at a final concentration of 2 µg/ml. Media was replenished every 2 days.

### DNA-vs RNA-based recovery

Approximately 500,000 HCT-116 cells were plated in a single well of a 6-well plate. Cells were transfected with 2.5 µg of the SP1-PBase plasmid (for a full list of plasmids, see Supp. Table 3) and 2.5 µg of the PB-SRT-Puro plasmid using Lipofectamine 3000 (Thermo Fisher #L3000015) following manufacturer’s instructions. After 24 hours, cells were split and plated 1:10 in each of three 10 cm dishes. Puromycin was then added and colonies were allowed to grow out under selection for two weeks. We obtained approximately 2,300 colonies. All cells were pooled together and split into two populations: one was subjected to DNA extraction, self-ligation, and inverse PCR, as described previously [35]; while the other underwent RNA extraction and SRT library preparation (see below).

### *In vitro* bulk calling card experiments

We cotransfected 10-12 replicates of HCT-116 cells with 5 µg of PB-SRT-Puro plasmid and 5 µg PBase plasmid via Neon electroporation (Thermo Fisher #MPK10025). Each replicate contained 2×10^6^ cells. As a negative control, we transfected one replicate of HCT-116 cells with 5 µg PB-SRT-Puro plasmid only. We used the following settings–pulse voltage: 1,530 V; pulse width: 20 ms; pulse number: 1. Each replicate was allowed to recover in a single well of a 6-well plate for 24 hours before being split 1:1 into a 10 cm dish and adding puromycin. Cells were grown under selection for one week, by which time almost all negative control transfectants were dead. We used the same experimental setup for experiments with PB-SRT-Puro and each of SP1-PBase, HyPBase, and SP1-HyPBase plasmids, as well as with SB-SRT-Puro and SB100X plasmids. Each replicate was cultured independently under aforementioned media conditions. After 7 days, we dissociated each replicate with trypsin-EDTA (Sigma #T4049) and created single cell suspensions in phosphate-buffered saline (PBS; Gibco #14190-136). Aliquots of each replicate were cryopreserved in cell culture media (see above) supplemented with 5% DMSO. The remaining cells were pelleted by centrifugation at 300g for 5 minutes. Cell pellets were either processed immediately or kept at −80°C in RNAProtect Cell Reagent (QIAGEN # 76526).

### Isolation of bulk RNA and reverse transcription

Total RNA was isolated from each replicate using the RNEasy Plus Mini Kit (QIAGEN #74134) following manufacturer’s instructions. Briefly, cell pellets were resuspended in 600 µl of Buffer RLT Plus with 1% 2-mercaptoethanol (Gibco #21985-023). Cells were homogenzied by vortexing. DNA was removed by running lysate through gDNA Eliminator spin columns, while RNA was bound by passing the flow-through over RNEasy spin columns. An on-column treatment with DNase (QIAGEN #79254) was also performed. After washing, RNA was eluted in 40 µl RNase-free H_2_O. RNA was quantitated using the Qubit RNA HS Assay Kit (Thermo Fisher #Q32852).

We performed first strand synthesis on each replicate with Maxima H Minus Reverse Transcriptase (Thermo Fisher #EP0752). We mixed 2 µg of total RNA with 1 µl 10 mM dNTPs (Clontech #639125) and 1 µl of 50 µM SMART_dT18VN primer (for a complete list of primer sequences, see Supp. Table 4), brought the total volume up to 14 µl, and incubated it at 65°C for 5 minutes. After transferring to ice and letting rest for 1 minute, we added 4 µl 5X Maxima RT Buffer, 1 µl RNaseOUT (Thermo Fisher #10777019), and 1 µl of 1:1 Maxima H Minus Reverse Transcriptase diluted in 1x RT Buffer (100 U). The solution was mixed by pipetting and incubated at 50°C for 1 hour followed by heat inactivation at 85°C for 10 minutes. Finally, we digested with 1 µl RNaseH (New England BioLabs #M0297S) at 37°C for 30 minutes. cDNA was stored at −20°C.

### Amplification of self-reporting transcripts from bulk RNA

The PCR conditions for amplifying self-reporting transcripts (i.e. transcripts derived from self-reporting transposons) involved mixing 1 µl cDNA template with 12.5 µl Kapa HiFi HotStart ReadyMix (Kapa Biosystems #KK2601), 0.5 µl 25 µM SMART primer, and either 1 µl of 25 µM SRT_PAC_F1 primer (in the case of puromycin selection) or 0.5 µl of 25 µM SRT_tdTomato_F1 primer (in the case of tdTomato screening). The mixture was brought up to 25 µl with ddH_2_O. Thermocycling parameters were as follows: 95°C for 3 minutes; 20 cycles of: 98°C for 20 seconds–65°C for 30 seconds–72°C for 5 minutes; 72°C for 10 minutes; hold at 4°C forever. As a control, cDNA quality can be assessed with exon-spanning primers for β-actin (see Supp. Table 4 for examples of human primers [95]) under the same thermocycling settings.

PCR products were purified using AMPure XP beads (Beckman Coulter #A63880). 12 µl of resuspended beads were added to the 25 µl PCR product and mixed homogenously by pipetting. After a 5-minute incubation at room temperature, the solution was placed on a magnetic rack for 2 minutes. The supernatant was aspirated and discarded. The pellet was washed twice with 200 µl of 70% ethanol (incubated for 30 seconds each time), discarding the supernatant each time. The pellet was left to dry at room temperature for 2 minutes. To elute, we added 20 µl ddH_2_O to the pellet, resuspended by pipetting, incubated at room temperature for 2 minutes, and placed on a magnetic rack for one minute. Once clear, the solution was transferred to a clean 1.5 ml tube. DNA concentration was measured on the Qubit 3.0 Fluorometer (Thermo Fisher #Q33216) using the dsDNA High Sensitivity Assay Kit (Thermo Fisher #Q32851).

### Generation of bulk RNA calling card libraries

Calling card libraries from bulk RNA were generated using the Nextera XT DNA Library Preparation Kit (Illumina #FC-131-1024). One nanogram of PCR product was resuspended in 5 µl ddH_2_O. To this mixture we added 10 µl Tagment DNA (TD) Buffer and 5 µl Amplicon Tagment Mix (ATM). After pipetting to mix, we incubated the solution in a thermocycler preheated to 55°C. The tagmentation reaction was halted by adding 5 µl Neutralization Tagment (NT) Buffer and was kept at room temperature for 5 minutes. The final PCR was set up by adding 15 µl Nextera PCR Mix (NPM), 8 µl ddH2O, 1 µl of 10 µM transposon primer (e.g. OM-PB-NNN) and 1 µl Nextera N7 indexed primer. The transposon primer anneals to the end of the transposon terminal repeat–*piggyBac,* in the case of OM-PB primers, or *Sleeping Beauty,* in the case of OM-SB primers–and contains a 3 base pair barcode sequence. Every N7 primer contains a unique index sequence that is demultiplexed by the sequencer. Each replicate was assigned a unique combination of barcoded transposon primer and indexed N7 primer, enabling precise identification of each library’s sequencing reads.

The final PCR was run under the following conditions: 95°C for 30 seconds; 13 cycles of: 95°C for 10 seconds–50°C for 30 seconds–72°C for 30 seconds; 72°C for 5 minutes; hold at 4°C forever. After PCR, the final library was purified using 30 µl (0.6x) AMPure XP beads, as described above. The library was eluted in 11 µl ddH_2_O and quantitated on an Agilent TapeStation 4200 System using the High Sensitivity D1000 ScreenTape (Agilent #5067-5584 and #5067-5585).

### Sequencing and analysis of bulk RNA calling card libraries

Multiple calling card libraries were pooled together for sequencing on the Illumina HiSeq 2500 platform. To increase the complexity of the library, PhiX was added at a final loading concentration of 50%. Reads were demultiplexed by the N7 index sequences added during the final PCR. Read 1 began with the 3 base pair barcode followed by the end of the transposon terminal repeat, culminating with the insertion site motif (TTAA in the case of *piggyBac*; TA in the case of *Sleeping Beauty*) before entering the genome. *piggyBac* reads were checked for exact matches to the barcode, transposon sequence, and insertion site at the beginning of reads before being hard trimmed using cutadapt [96] with the following settings: -g “^NNNTTTACGCAGACTATCTTTCTAGGGTTAA” --minimum-length 1 --discard-untrimmed -e 0 --no-indels, where NNN is replaced with the primer barcode. *Sleeping Beauty* libraries were trimmed with the following settings: -g “^NNNTAAGTGTATGTAAACTTCCGACTTCAACTGTA” --minimum-length 1 --discard-untrimmed -e 0 --no-indels. Reads passing this filter were then trimmed of any trailing Nextera adapter sequence, again using cutadapt and the following settings: -a “CTGTCTCTTATACACATCTCCGAGCCCACGAGACTNNNNNNNNNNTCTCGTATGCCGTCTTCTGCTTG” -- minimum-length 1. The remaining reads were aligned to the human genome (build hg38) with Novoalign 3 (Novocraft Technologies) and the following settings: -n 40 -o SAM -o SoftClip. Aligned reads were validated by confirming that they mapped adjacent to the insertion site motif. Successful reads were then converted to calling card format (.ccf; see http://wiki.wubrowse.org/Calling_card) and visualized on the WashU Epigenome Browser v46 (http://epigenomegateway.wustl.edu/legacy/).

### *In vitro* single cell calling card experiments

N2a and K562 cells were cultured and transfected identically as HCT-116 cells, with the following exceptions: K562 cells were grown in RPMI 1640 Medium (Gibco #11875-085); for K562 cells, Neon electroporation settings were–pulse voltage: 1,450 V; pulse width: 10 ms; pulse number: 3; for N2a cells, Neon electroporation settings were–pulse voltage: 1,050 V; pulse width: 30 ms; pulse number: 2. For N2a cells, one replicate (2×10^6^ cells) was transfected with 5 µg PB-SRT-Puro and 5 µg HyPBase, while another replicate was transfected with 5 µg PB-SRT-Puro only. For K562 cells, 4 replicates received both plasmids and one received the SRT alone. After 1 week of selection, N2a or K562 cells were mixed with transfected HCT-116 cells and then underwent single cell RNA-seq library preparation. For the species mixing experiment, cells were classified as either human or mouse if at least 80% of self-reporting transcripts in that cell mapped to the human or mouse genome, respectively, and as a multiplet. The estimated multiplet rate was calculated by doubling the observed percentage of human-mouse multiplet, to account for human-human and mouse-mouse doublets.

### Single cell RNA-seq library preparation

Single cell RNA-seq libraries were prepared using 10x Genomics’ Chromium Single Cell 3’ Library and Gel Bead Kit (v2 chemistry; #120267). Each replicate was targeted for recovery of 6,000 cells. Library preparation followed a modified version of the manufacturer’s protocol. We prepared the Single Cell Master Mix without RT Primer, replacing it with an equivalent volume of Low TE Buffer. GEM generation and GEM-RT incubation proceeded as instructed. At the end of Post GEM-RT cleanup, we added 36.5 µl Elution Solution I and transferred 36 µl of the eluted sample to a new tube (instead of 35.5 µl and 35 µl, respectively). The eluate was split into two 18 µl aliquots and kept at –20°C until ready for further processing. One fraction was kept for single cell calling cards library preparation (see next section), while the other half was further processed into a single cell RNA-seq library.

We then added the RT Primer sequence to the products in the scRNA-seq aliquot. We created an RT master mix by adding 20 µl of Maxima 5X RT Buffer, 20 µl of 20% w/v Ficoll PM-400 (GE Healthcare #17030010), 10 µl of 10 mM dNTPs (Clontech #639125), 2.5 µl RNase Inhibitor (Lucigen), and 2.5 µl of 100 µM 10x_TSO. To this solution we added 18 µl of the first RT product and 22 µl of ddH_2_O. Finally, we added 5 µl Maxima H Minus Reverse Transcriptase, mixed by flicking, and centrifuged briefly. This reaction was incubated at 25°C for 30 minutes followed by 50°C for 90 minutes and heat inactivated at 85°C for 5 minutes.

The solution was purified using DynaBeads MyOne Silane (Thermo Fisher #37002D) following 10x Genomics’ instructions, beginning at “Post GEM-RT Cleanup – Silane DynaBeads” step D. The remainder of the single cell RNA-seq protocol, including purification, amplification, fragmentation, and final library amplification, followed manufacturer’s instructions.

### Single cell calling cards library preparation

To amplify self-reporting transcripts from single cell RNA-seq libraries, we took 9 µl of RT product (the other half was kept in reserve) and added it to 25 µl Kapa HiFi HotStart ReadyMix and 15 µl ddH_2_O. We then prepared a PCR primer cocktail comprising 5 µl of 100 µM Bio_Illumina_Seq1_scCC_10X_3xPT primer, 5 µl of 100 µM Bio_Long_PB_LTR_3xPT, and 10 µl of 10 mM Tris-HCl, 0.1 mM EDTA buffer (IDT #11-05-01-13). One µl of this cocktail was added to the PCR mixture and placed in a thermocycler (Eppendorf MasterCycler Pro). Thermocycling settings were as follows: 98°C for 3 minutes; 20-22 cycles of 98°C for 20 seconds–67°C for 30 seconds–72°C for 5 minutes; 72°C for 10 minutes; 4°C forever. PCR purification was performed with 30 µl AMPure XP beads (0.6x ratio) as described previously. The resulting library was quantitated on an Agilent TapeStation 4200 System using the High Sensitivity D5000 ScreenTape (Agilent #5067-5592 and #5067-5593).

Single cell calling card library preparation was performed using the Nextera Mate Pair Sample Prep Kit (Illumina #FC-132-1001) with modifications to the manufacturer’s protocol. The library was circularized by bringing 300 fmol (approximately 200 ng) of DNA up to a final volume of 268 µl with ddH_2_O, then adding 30 µl Circularization Buffer 10x and 2 µl Circularization Ligase (final concentration: 1 nM). This reaction was incubated overnight (12-16 hours) at 30°C. After removal of linear DNA (following manufacturer’s instructions), we sheared the library on a Covaris E220 Focused-ultrasonicator with the following settings–peak power intensity: 200; duty factor: 20%; cycles per burst: 200; time: 40 seconds; temperature: 6°C.

The library preparation proceeded per manufacturer’s instructions until adapter ligation. We designed custom adapters (Supp. Table 4) so that the standard Illumina sequencing primers would not interfere with our library. Adapters were prepared by combining 4.5 µl of 100 µM scCC_P5_adapter, 4.5 µl of 100 µM scCC_P7_adapter, and 1 µl of NEBuffer 2 (New England BioLabs #B7002S), then heating in a thermocycler at 95°C for 5 minutes, then holding at 70°C for 15 minutes, then ramping down at 1% until it reached 25°C, holding at that temperature for 5 minutes, before keeping at 4°C forever. One microliter of this custom adapter mix was used in place of the manufacturer’s recommended DNA Adapter Index.

The ligation product was cleaned per manufacturer’s instructions. For the final PCR, the master mix was created by combining 20 µl Enhanced PCR Mix with 28 µl of ddH_2_O and 1 µl each of 25 µM scCC_P5_primer and 25 µM scCC_P7_primer. This was then added to the streptavidin bead-bound DNA and amplified under the following conditions: 98°C for 30 seconds; 15 cycles of: 98°C for 10 seconds– 60°C for 30 seconds–72°C for 2 minutes; 72°C for 5 minutes; 4°C forever. All of the PCR supernatant was transferred to a new tube and purified with 35 µl (0.7x) AMPure XP beads following manufacturer’s instructions. The final library was eluted in 25 µl Elution Buffer (QIAGEN #19086) and quantitated on an Agilent TapeStation 4200 System using the High Sensitivity D1000 ScreenTape.

### Sequencing and analysis of scRNA-seq libraries

scRNA-seq libraries were sequenced on either Illumina HiSeq 2500 or NovaSeq S1 machines. Reads were analyzed using 10x Genomics’ cellranger 2.1.0 with the following settings: --expect-cells=6000 -- chemistry=SC3Pv2 --localcores=16 --localmem=30. The digital gene expression matrices from 10x were then further processed with scanpy 1.3.7 [97] for identification of highly variable genes, dimensionality reduction, and Louvain clustering. We cross-referenced gene expression data with published datasets [98] to assign cell types. The species-mixing analysis was performed using Drop-seq_tools 1.11 [59].

### Sequencing and analysis of scCC libraries

scCC libraries were sequenced on Illumina NextSeq 500 machines (v2 Reagent Cartridges) with 50% PhiX. We used the standard Illumina primers for read 1 and index 2 (BP10 and BP14, respectively), and custom primers for read 2 and index 1 (Supp. Table 4). Read 1 sequenced the cell barcode and unique molecular index of each self-reporting transcript. Read 2 began with GGTTAA (end of the *piggyBac* terminal repeat and insertion site motif) before continuing into the genome. Reads containing this exact hexamer were trimmed using cutadapt with the following settings: -g “^GGTTAA” --minimum-length 1 -- discard-untrimmed -e 0 --no-indels. Reads passing this filter were then trimmed of any trailing P7 adapter sequence, again using cutadapt and with the following settings: -a “AGAGACTGGCAAGTACACGTCGCACTCACCATGANNNNNNNNNATCTCGTATGCCGTCTTCTGCTTG” -- minimum-length 1. Reads passing these filters were aligned using 10x Genomics’ cellranger with the following settings: --expect-cells=6000 --nosecondary --chemistry=SC3Pv2 --localcores=16 -- localmem=30. This workflow also managed barcode validation and collapsing of UMIs. Aligned reads were validated by verifying that they mapped adjacent to TTAA tetramers. Reads were then converted to calling card format (.ccf, see above). Finally, to minimize the presence of intermolecular artifacts, we required that each insertion must have been tagged by at least two different UMIs. We used the set of validated cell barcodes from each scRNA-seq library to demultiplex library-specific barcoded insertions from the scCC data. This approach requires no shared cell barcodes between scCC (and scRNA-seq) libraries. As a result, we excluded insertions from non-unique cell barcodes, which represented a very small number of total cells lost (< 1% per multiplexed library).

### Peak calling

We called peaks in calling card data using Bayesian blocks [99], a noise-tolerant algorithm for segmenting discrete, one-dimensional data, using the astroML 0.3 implementation [100]. Bayesian blocks segments the genome into non-overlapping blocks where the density of calling card insertions is uniform. By comparing the segmentation against a background model, we were able to use Poisson statistics to assess whether a given block shows statistically significant enrichment for insertions. Let *B* = {b_1_, b_2_, … b_n_} represent the set of blocks found by performing Bayesian block segmentation on all insertions from a TF-directed experiment (e.g. SP1-PBase). For each block *b_i_*, let *x_i_* be the number of insertions in that block in the TF-directed experiment. Similarly, let 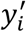 be the number of insertions in that block in the undirected experiment (e.g. PBase) normalized to the total number of insertions found in the TF-directed experiment. Then, for each block we calculated the Poisson *p-*value of observing at least *x_i_* insertions assuming a Poisson distribution with expectation 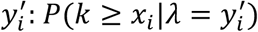. We accepted all blocks that were significant beyond a particular *p*-value threshold.

For bulk analysis of SP1-PBase and SP1-HyPBase insertions, we added a pseudocount of 0.1 to all blocks and used *p*-value cutoffs of 10^−6^ and 10^−22^, respectively. For single cell analysis of SP1-HyPBase insertions, we added a pseudocount of 1 to all blocks and used a p-value cutoff of 10^−9^. All three of these values were beyond a Bonferroni-corrected 6 of 0.05. We polished peak calls by merging statistically-significant blocks that were within 250 bases of each other and by aligning block edges to coincide with TTAAs.

To identify BRD4 binding sites from undirected *piggyBac* insertions, we segmented those insertions using Bayesian blocks. For each block *b_i_*, we let *x_i_* denote the number of undirected insertions in that block. We also calculated 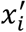, the expected number of insertions in block *b_i_* assuming *piggyBac* insertions were distributed uniformly across the genome. We did this by dividing the total number of mappable TTAAs in the genome by the total number of undirected insertions, then multiplying this value by the number of mappable TTAAs in block *b_i_*. Then, for each block we calculated the Poisson 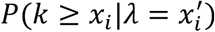. We accepted all blocks that were significant beyond a particular *p*-value threshold. Finally, we merged statistically-significant blocks that were within 12,500 bases of each other [51,57].

For the bulk PBase and HyPBase analysis, we used *p*-value cutoffs of 10^−30^ and 10^−62^, respectively. For both *in vitro* and *in vivo* single cell HyPBase analyses, we used a *p-*value cutoff of 10^−9^. To call differentially-bound loci between upper and lower cortical layer neurons, we used the same framework as described above for SP1 but did reciprocal enrichment analyses where the upper layer insertions were used as the “experiment” track and the lower layer insertions were used as the “control” track, and vice-versa. Here again we used a *p*-value cutoff of 10^−9^.

Density tracks were generated by taking the Bayesian blocks segmentation of each calling card dataset and, for each block, calculating the normalized number of insertions (insertions per million mapped insertions, or IPM) and dividing by the length of the block in kilobases. This was plotted as a bedgraph file with smoothing applied in the WashU Epigenome Browser (25 pixel windows).

### SP1 binding analysis in HCT-116 cells

We compared our SP1 peak calls to a publicly-available ChIP-seq dataset [54] as well as an input control file (Supp. Table 5). See below for more details on aligning and analyzing ChIP-seq data. We collated a list of unique TSSs by taking the 5’-most coordinates of RefSeq Curated genes in the hg38 build (UCSC Genome Browser). A list of CpG islands in HCT-116 cells and their methylation statuses were derived from previously-published Methyl-seq data [101]. We used the liftOver tool (UCSC) to convert coordinates from hg18 to hg38. We tested for enrichment in SP1-directed insertions at TSSs, CpG islands, and unmethylated CpGs with the *G* test of independence. For motif discovery we used MEME-ChIP 4.11.2 [102] with a dinucleotide shuffled control and the following settings: -dna -nmeme 600 -seed 0 -ccut 250 - meme-mod zoops -meme-minw 4 -meme-nmotifs 5.

### BRD4 sensitivity, specificity, and precision analysis in HCT-116 cells

We used a published BRD4 ChIP-seq dataset [53] to identify BRD4-bound super-enhancers in HCT-116 cells, following previously-described methods [51,52]. We first called peaks using MACS 1.4.1 [103] at *p* < 10^−9^, then fed this list into ROSE 0.1 (http://younglab.wi.mit.edu/super_enhancer_code.html). We discarded artifactual loci less than 2,000 bp in size, yielding a final list of 162 super-enhancers. To evaluate sensitivity, we used bedtools 2.27.1 [104] to ask what fraction of *piggyBac* peaks, at various *p*-value thresholds, overlapped the set of BRD4-bound super-enhancers. To measure specificity, we created a list of regions predicted to be insignificantly enriched (*p* > 0.1) for BRD4 ChIP-seq signal. We then sampled bases from this region such that the distribution of peak sizes was identical to that of the 162 super-enhancers. We sampled to 642x coverage, sufficient to cover each base with one peak, on average. We then asked what fraction of our *piggyBac* peaks overlapped these negative peaks and subtracted that value from 1 to obtain specificity. Finally, we calculated precision, or positive predictive value, by dividing the total number of detected super-enhancer peaks by the sum of the super-enhancer peaks and the false positive peaks.

### Downsampling and replication analysis

When performing downsampling analyses on calling card insertions, we randomly sampled insertions without replacement and in proportion to the number of reads supporting each insertion. Peaks were called on the downsampled insertions at a range of *p*-value cutoffs. Linear interpolation was performed using numpy 1.15 and visualized using matplotlib 3.0. Replication was assessed by splitting calling card insertions into two, approximately equal, files based on their barcode sequences. Each new file was treated as a single biological experiment. For each peak called from the joint set of all insertions, we plotted the number of normalized insertions (IPM) in one replicate on the *x*-axis and the other replicate on *y*-axis.

### ChIP-seq and chromatin state analyses

We aligned raw reads using Novoalign with the following settings for single-end datasets: -o SAM -o SoftClip, while paired-end datasets were mapped with the additional flag -i PE 200-500. To calculate and visualize the fold enrichment in ChIP-seq signal at calling card peaks, we used deeptools 3.0.1 [105]. We tested for significant mean enrichment in BRD4 ChIP-seq signal at *piggyBac* peaks over randomly shuffled control peaks with the Kolmogorov-Smirnov test. Chromatin state analysis was performed using ChromHMM 1.15 as previously described [106]. For each chromatin state, we plotted the mean and standard deviation of the rate of normalized insertions per kilobase (IPM/kb).

### SRT-tdTomato fluorescence validation

To test the fluorescence properties of the SRT-tdTomato construct, we transfected K562 cells as previously described with either 1 µg of pUC19 plasmid (New England BioLabs #N3041S); 0.5 µg of PB-SRT-tdTomato plasmid and 0.5 µg pUC19; 0.5 µg of PB-SRT-tdTomato and 0.5 µg PBase plasmid; and 0.5 µg of PB-SRT-tdTomato and 0.5 µg HyPBase plasmid. Cells were allowed to expand for 8 days, after which fluorescence activity was assayed on an Attune NxT Flow Cytometer (Thermo Fisher) with an excitation wavelength of 561 nm. Flow cytometery data were visualized using FlowCal 1.2.0 [107]. We also performed bulk RNA calling cards on HEK293T cells transfected with SRT-tdTomato with or without HyPBase plasmid. While these cells were not sorted based on fluorescence activity, the SRT library from cells transfected with both SRT and transposase were more complex and contained many more insertions than the library from cells receiving SRT alone (Supp. Fig. 1A).

### *In vivo* single cell calling cards experiments

All mouse experiments were done following procedures described in [66]. In brief, we cloned the PB-SRT-tdTomato and HyPBase constructs into AAV vectors. The Hope Center Viral Vectors Core at Washington University in St. Louis packaged each construct in AAV9 capsids. Titers for each virus ranged between 1.1×10^13^ and 2.2×10^13^ viral genomes/ml. We mixed equal volumes of each virus and performed intracranial cortical injections of the mixture into newborn wild-type C57BL/6J pups (P0-2). As a gating control, we injected one litter-matched animal with AAV9-PB-SRT-tdTomato only. After 2 to 4 weeks, we sacrificed mice and dissected the cortex (8 libraries) or hippocampus (1 library). All animal practices and procedures were approved by the Washington University in St. Louis Institutional Animal Care and Use Committee (IACUC) in accordance with National Institutes of Health (NIH) guidelines.

Tissues were dissociated to single suspensions following a modification of previously published methods [108,109]. We incubated samples in a papain solution containing Hibernate-A (Gibco #A1247501) with 5% v/v trehalose (Sigma-Aldrich #T9531), 1x B-27 Supplement (Gibco #17504044), 0.7 mM EDTA (Corning #36-034-Cl), 70 µM 2-mercaptoethanol (Gibco #21985023), and 2.8 mg/ml papain (Worthington Chemical Corporation #LS003118). After incubation at 37°C, cells were treated with DNaseI (Worthington Chemical Corporation #NC9924263), triturated through increasingly narrow fire-polished pipettes, and passed through a 40-micron filter prewetted with resuspension solution: Hibernate-A containing 5% v/v trehalose, 0.5% Ovomucoid Trypsin Inhibitor (Worthington Chemical Corporation #NC9931428), 0.5% Bovine Serum Albumin (BSA; Sigma-Aldrich #A9418), 33 µg/ml DNaseI, and 1x B-27 Supplement. The filter was washed with 6 ml of resuspension solution. The resulting suspension was centrifuged for 4 minutes at 250 g. The supernatant was discarded. The pellet was then resuspended in 2 ml of resuspension solution and resuspended by gentle pipetting.

We eliminated subcellular debris using gradient centrifugation. We first prepared a working solution of 30% w/v OptiPrep Density Gradient Medium (Sigma-Aldrich #D1556) mixed with an equal volume of 1x Hank’s Balanced Salt Solution (HBSS; Gibco #14185052) with 0.5% BSA. We then prepared solutions of densities 1.057, 1.043, 1.036, and 1.029 g/ml using by combining the working solution with resuspension solution at ratios of 0.33:0.67, 0.23:0.77, 0.18:0.82, and 0.13:0.87, respectively. We layered 1 ml aliquots of each solution in a 15 ml conical tube beginning with the densest solution on the bottom. The cell suspension was added last to the tube and centrifuged for 20 minutes at 800 g at 12°C. The top layer was then aspirated and purified cells were isolated from the remaining layers. These cells were then resuspended in FACS buffer: 1x HBSS, 2 mM MgCl_2_ (Sigma-Aldrich #M4880), 2 mM MgSO_4_ (Sigma-Aldrich #M2643), 1.25 mM CaCl_2_ (Sigma-Aldrich #C7902), 1 mM D-glucose (Sigma-Aldrich #G7021), 0.02% BSA, and 5% v/v trehalose. Cells were centrifuged for 4 minutes at 250 g, the supernatant was discarded, and the pellet was resuspended in FACS buffer by gentle pipetting.

Cells were then sorted based on fluorescence activity. As a gating control, we analyzed cells from cortices injected with AAV9-PB-SRT-tdTomato only. We then collected cells from brains transfected with AAV9-PB-SRT-tdTomato and AAV9-HyPBase whose fluorescence values exceeded the gate. After sorting, cells were centrifuged for 3 minutes at 250 g. The supernatant was discarded and cells were resuspended in FACS buffer at a concentration appropriate for 10x Chromium 3’ scRNA-seq library preparation.

### *In vivo* single cell calling cards analysis and validation

Single cell RNA-seq and single cell calling card libraries were prepared, sequenced, and analyzed as described above. Cell types were assigned based on the expression of key marker genes and cross-referenced with recent cortical scRNA-seq datasets [62-65]. Brd4-bound peak calls were validated by comparing to a previously published cortical H3K27ac ChIP-seq dataset [68] (Supp. Table 5). Read alignment and statistical analysis were performed as described above.

The specificity of Brd4-bound gene expression in astrocytes and neurons was analyzed by first identifying all genes within 10,000 bases of astrocyte and neuronal Brd4 peaks. Although assigning an enhancer to its target gene is a difficult problem, using the nearest gene is common practice [110]. To control for sensitivity of gene detection, we downsampled the neuron insertions to the same number of astrocyte insertions, then called peaks and identified nearby genes in this subset. We used gene expression data from a bulk RNA-seq dataset [69] to compute the specificity of gene expression between astrocytes and neurons. We first discarded genes whose expression was not measured, and then set the value for genes with 0.1 FPKM to zero (to better distinguish non-expressed genes from lowly-expressed genes). Finally, for each gene 7, we calculated the specificity as 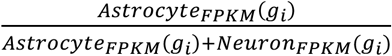. Thus, a value of 0 denotes a gene purely expressed in neurons, a value of 0.5 for a gene equally expressed in both cell types, and a value of 1 for a gene purely expressed in astrocytes. We plotted distributions of gene expression specificity for the set of astrocyte-bound genes and the downsampled astrocyte-bound genes. Gene Ontology analysis was performed on the same sets of genes using PANTHER 14.0 [70] on the “GO biological process complete” database. Fisher’s exact test was used to compute *p*-values, which were then subject to Bonferroni correction.

