## Supplemental Materials for "Self-reporting transposons enable simultaneous readout of gene expression and transcription factor binding in single cells"

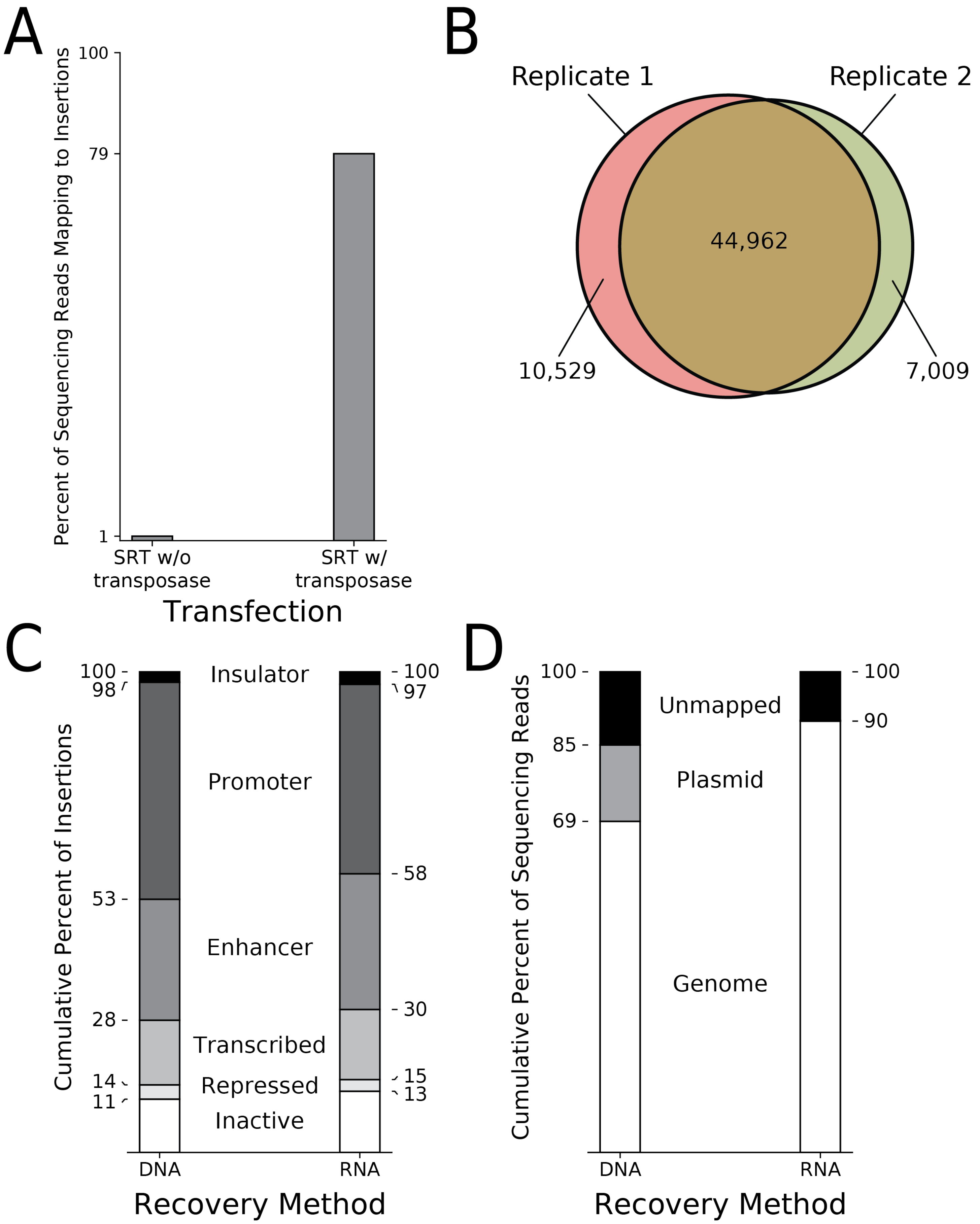
A
Percent of Sequencing Reads Mapping to Insertions
100
79
1
SRT w/o transposase
SRT w/ transposase
Transfection
B
Replicate 1
Replicate 2
44,962
10,529
7,009
C
Cumulative Percent of Insertions
100
98
53
28
14
11
100
97
58
30
15
13
Insulator
Promoter
Enhancer
Transcribed
Repressed
Inactive
DNA
RNA
Recovery Method
D
Cumulative Percent of Sequencing Reads
100
85
69
100
90
Unmapped
Plasmid
Genome
DNA
RNA
Recovery Method

**Supplemental Figure 1: Properties of self-reporting transposons.** (A) Analysis of bulk RNA calling card libraries prepared from HEK293T cells transfected with PB-SRT-tdTomato with and without HyPBase transposase. The transposase is required for efficient and complex library generation. (B) Technical replication of bulk RNA calling cards from HCT-116 cells transfected with PB-SRT-Puro and SP1-PBase. Over 80% of insertions in each trial were shared between both replicates. (C) No significant differences were observed between DNA- and RNA-based recovery of SP1-directed insertions with respect to chromatin state in HCT-116 cells. (D) The self-cleaving ribozyme eliminates recovery of un-excised transposons when calling card libraries are prepared from RNA but not DNA.

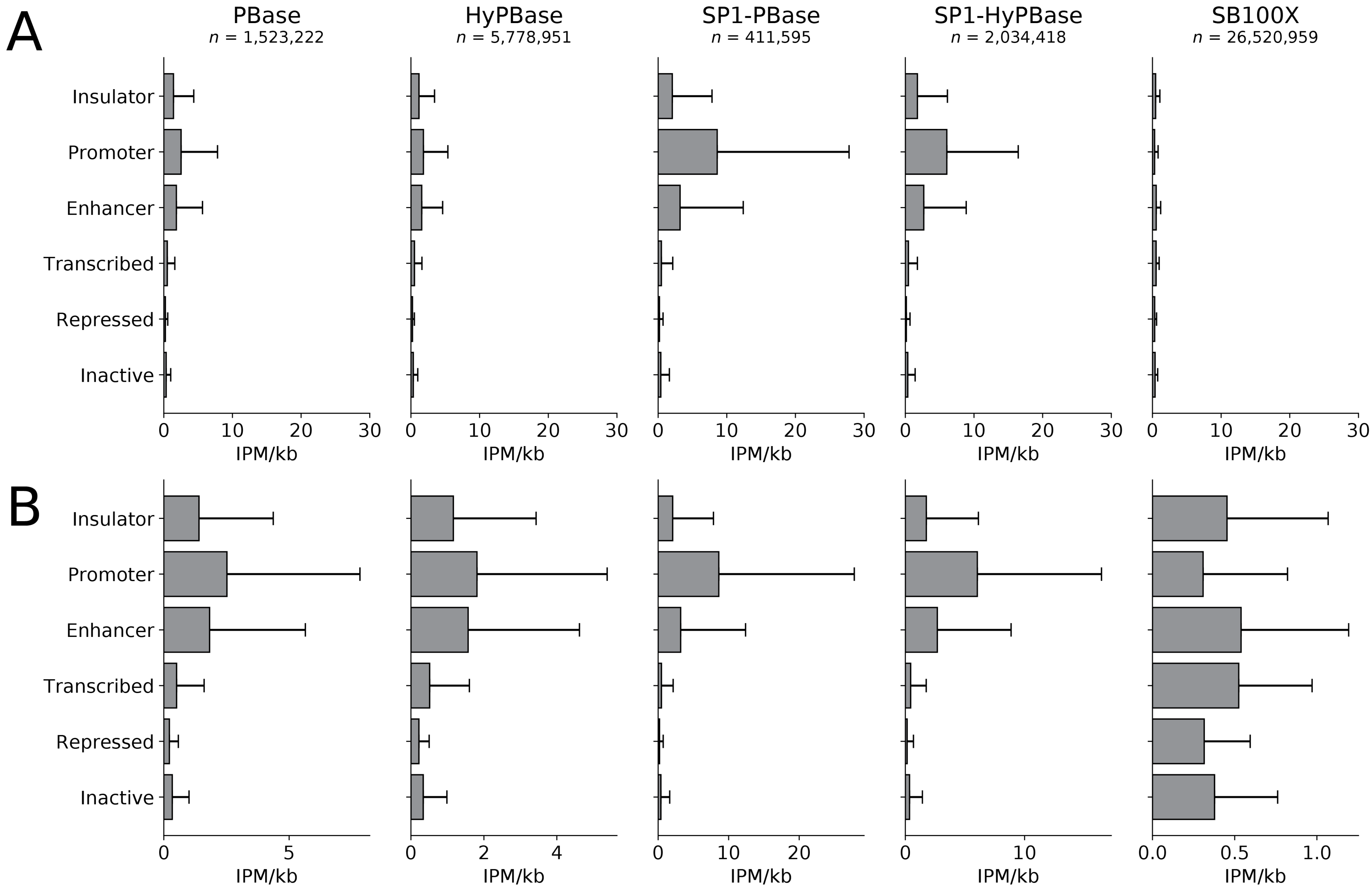


**Supplemental Figure 2: *piggyBac*, SP1-*piggyBac* fusions, and *Sleeping Beauty* display different local transposition rates depending on chromatin state.** (A) Chromatin state analysis on local rates of transposition. Undirected and SP1-directed *piggyBac* transposases show different preferences for chromatin states. Undirected *piggyBac* favors promoters and enhancers, while SP1-*piggyBac* fusions show marked preference for promoters. *Sleeping Beauty* shows uniform distribution of insertions across all chromatin states. Bars represent means; error bars are standard deviations. (B) Same data as (A) but with different *x*-axes for each graph. IPM: insertions per million mapped insertions; kb: kilobase.

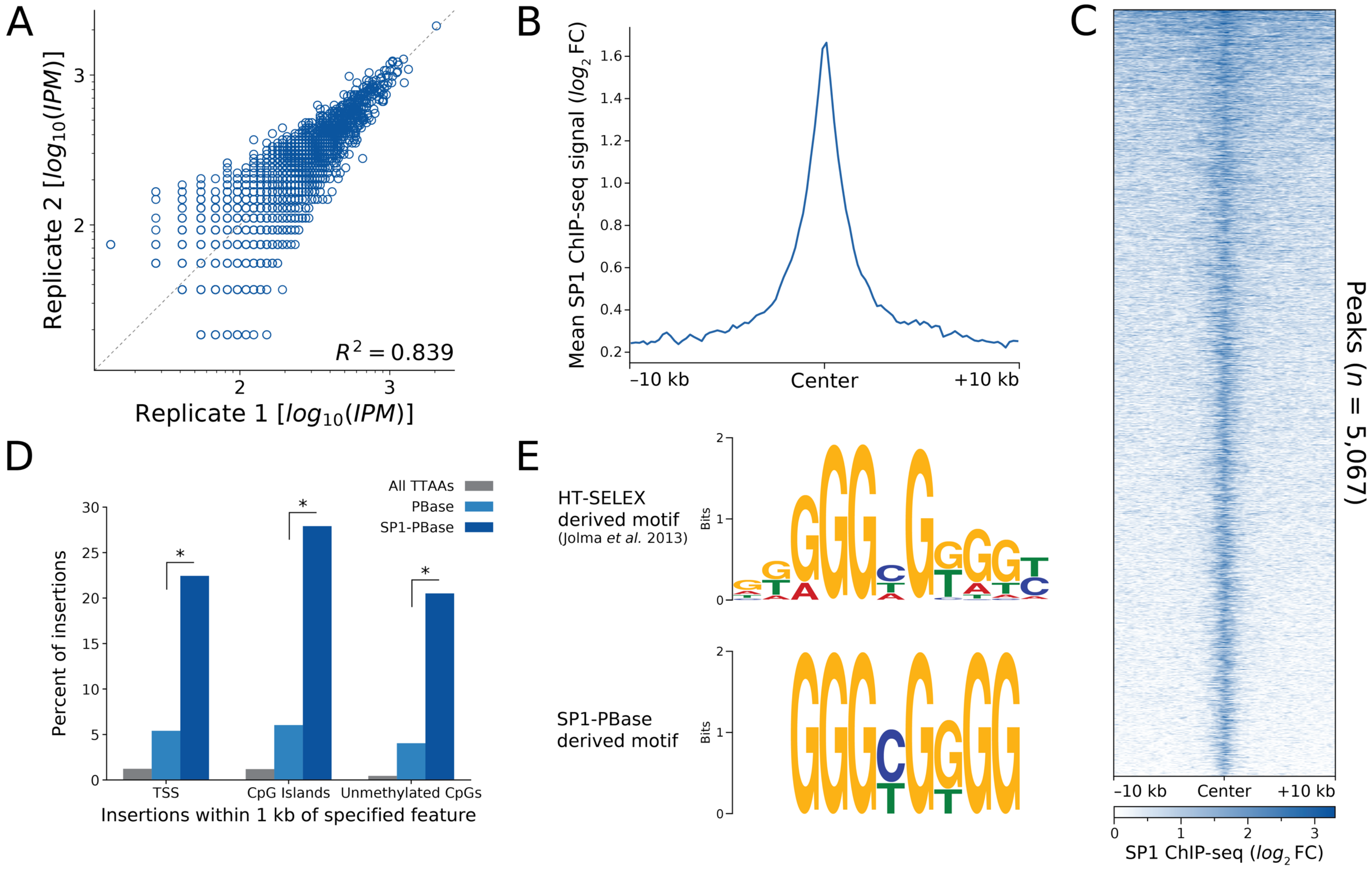


**Supplemental Figure 3: SP1 fused to *piggyBac* redirects insertions to SP1 binding sites.** (A) SP1 peaks show high reproducibility between biological replicates. Each circle represents a peak; *x* and *y* coordinates represent normalized insertions in each biological replicate. (B) Mean SP1 ChIP-seq profile across all SP1 peaks shows strong central enrichment. (C) Heatmap of SP1 ChIP-seq signal across all SP1 peaks, expressed as $log_2$(FC) over the input control. (D) SP1-PBase shows enrichment of insertions to transcription start sites (TSSs), CpG islands, and unmethylated CpGs, all known biological targets of SP1. Each enrichment was statistically significant at $p < 10^{-9}$ ($G$ test of independence). (E) Motif discovery performed on SP1 peaks shows good concordance with an orthogonally-derived SP1 motif. IPM: insertions per million mapped insertions; FC: fold change.

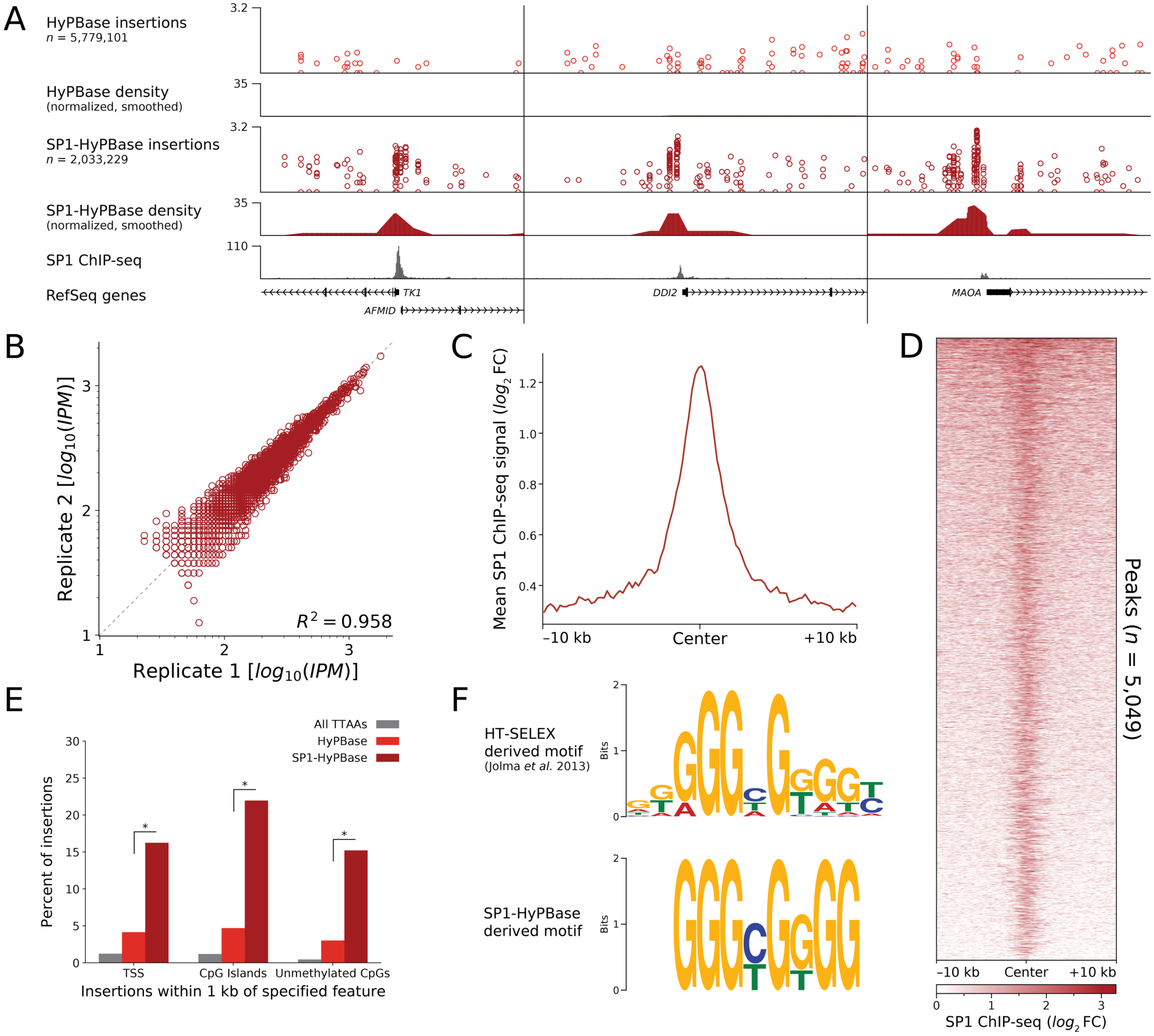


**Supplemental Figure 4: SP1 fused to hyperactive *piggyBac* (SP1-HyPBase) also redirects insertions to SP1 binding sites.** (A) SP1-HyPBase, like SP1-PBase, can also be used to identify SP1 binding sites. (B) Insertions at SP1-HyPBase-derived peaks show high reproducibility between biological replicates. (C) Mean SP1 ChIP-seq profile at peaks shows strong central enrichment. (D) Heatmap of SP1 ChIP-seq signal across all peaks, expressed as $log_2$(FC) over the input control. (E) SP1-HyPBase redirects insertions to TSSs, CpG islands, and unmethylated CpGs ($p < 10^{-9}$, $G$ test of independence). (F) Motif analysis of SP1-HyPBase peaks identifies the SP1 motif. IPM: insertions per million mapped insertions; FC: fold change.

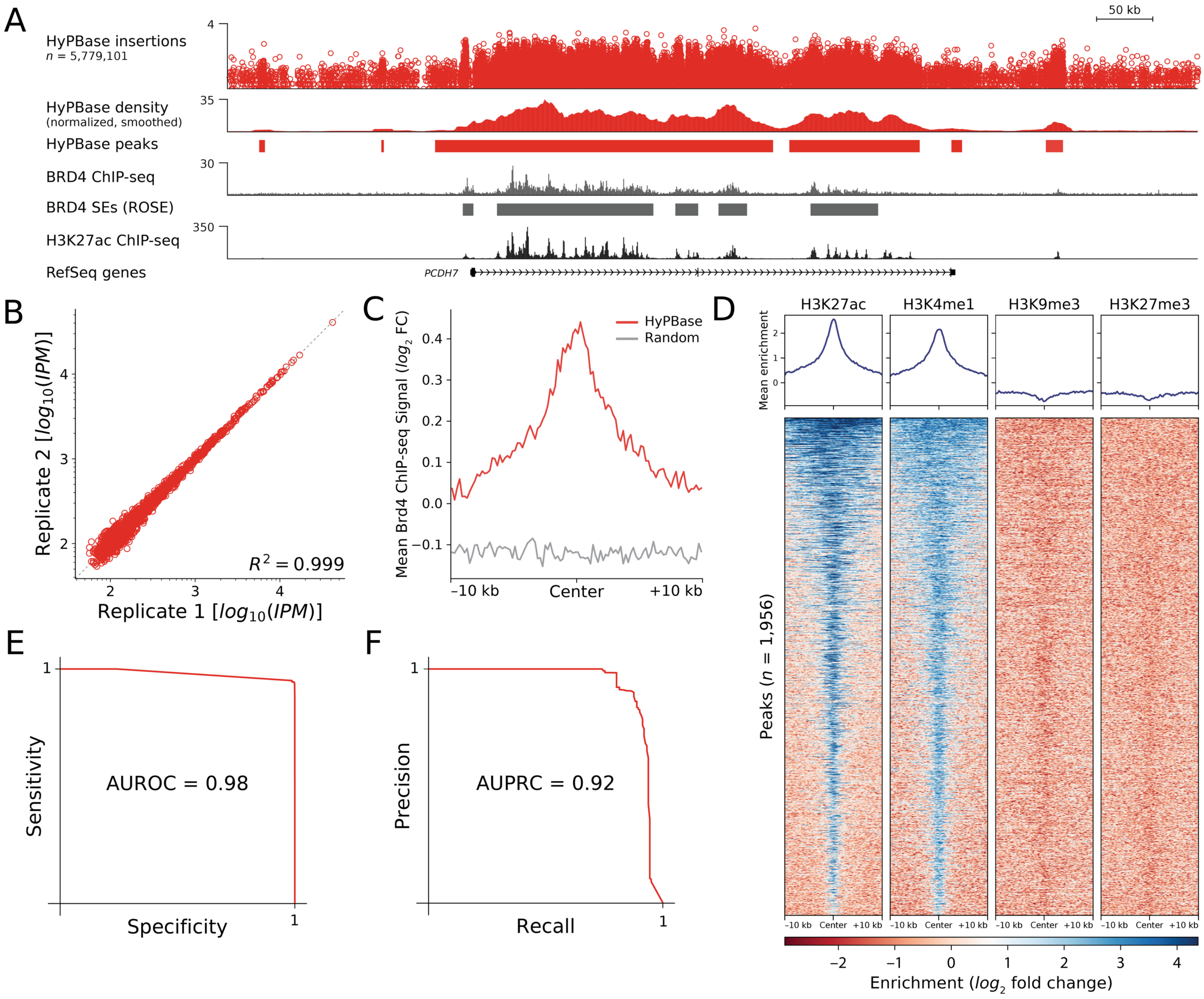
A
50 kb
HyPBase insertions
n = 5,779,101
4
HyPBase density
(normalized, smoothed)
35
HyPBase peaks
BRD4 ChIP-seq
30
BRD4 SEs (ROSE)
H3K27ac ChIP-seq
350
RefSeq genes
PCDH7
B
Replicate 2 [log10(IPM)]
Replicate 1 [log10(IPM)]
R² = 0.999
C
Mean Brd4 ChIP-seq Signal (log2 FC)
HyPBase
Random
−10 kb
Center
+10 kb
D
H3K27ac
H3K4me1
H3K9me3
H3K27me3
Mean enrichment
Peaks (n = 1,956)
−10 kb Center +10 kb
Enrichment (log2 fold change)
E
Sensitivity
Specificity
AUROC = 0.98
F
Precision
Recall
AUPRC = 0.92

**Supplemental Figure 5: Undirected hyperactive *piggyBac* (HyPBase) insertions also mark Brd4-bound super-enhancers.** (A) Undirected HyPBase, like PBase, shows non-uniform densities of insertions in BRD4-bound regions. (B) Densities of insertions are reproducible at HyPBase peaks. (C) Mean BRD4 ChIP-seq profile at HyPBase peaks compared to randomly chosen peaks. The BRD4 enrichment is significant with $p < 10^{-9}$ (KS test). (D) Undirected HyPBase peaks are strongly correlated with H3K27ac and H3K4me1 and mildly anti-correlated with H3K9me and H3K27me3, consistent with these regions being enhancers. (E) Receiver-operator characteristic curve for detecting BRD4-bound SEs with undirected HyPBase peaks. (F) Precision-recall curve for detecting SEs with undirected HyPBase peaks. IPM: insertions per million mapped insertions; KS: Kolmogorov-Smirnov; FC: fold change.

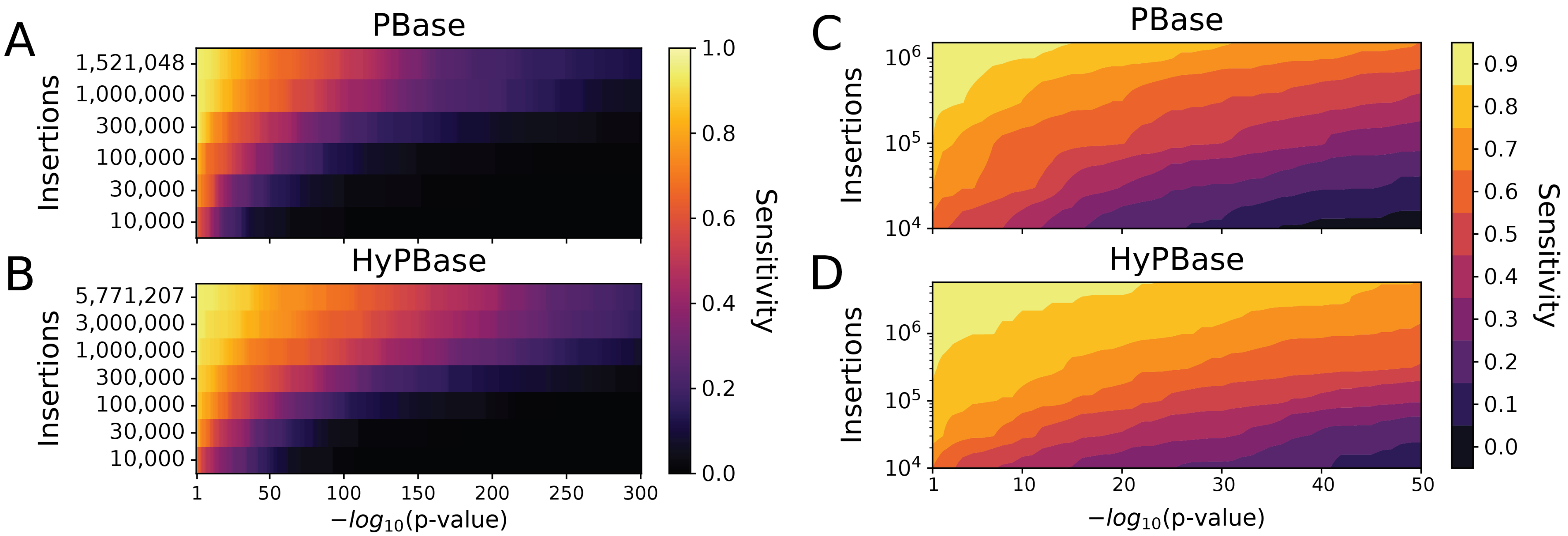


**Supplemental Figure 6: Downsampling PBase and HyPBase insertions affects sensitivity to BRD4-bound super-enhancers.** (A) Downsampling analysis of BRD4-bound SE detection by PBase insertions at various *p*-value thresholds. (B) Downsampling analysis applied to HyPBase insertions. (C) Linear interpolation applied to (A) to predict SE sensitivity across a range of insertions. (D) Linear interpolation applied to (B).

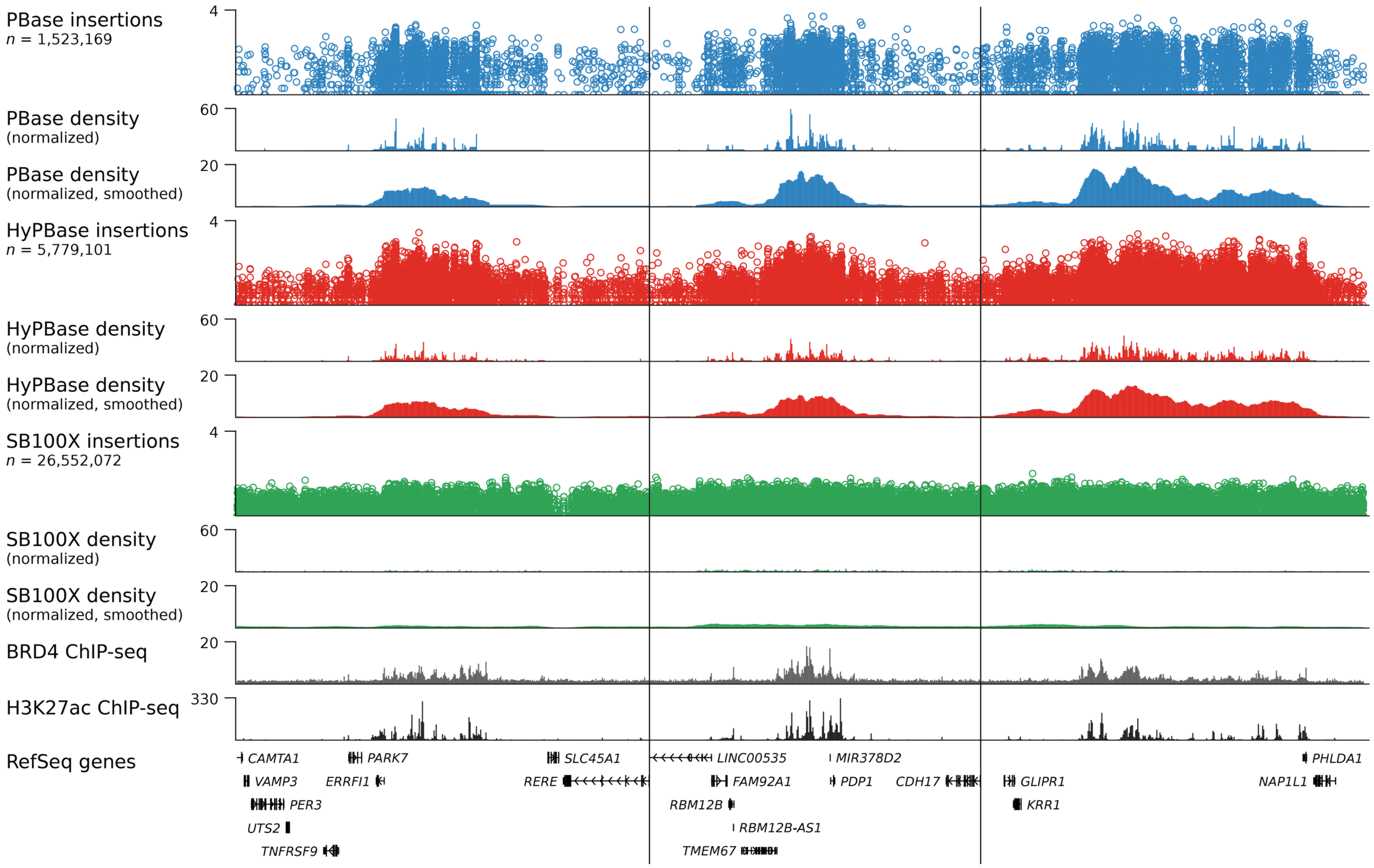


**Supplemental Figure 7: Examples of BRD4-bound super-enhancers identified by PBase and HyPBase calling cards.** Three different loci exhibiting non-uniform densitites of *piggyBac* insertions. These densities correlate well with BRD4 and H3K27ac ChIP-seq data. Density tracks are shown before and after smoothing. *Sleeping Beauty* does not show the same preference for BRD4-bound regions as *piggyBac* but instead appears uniformly distributed.

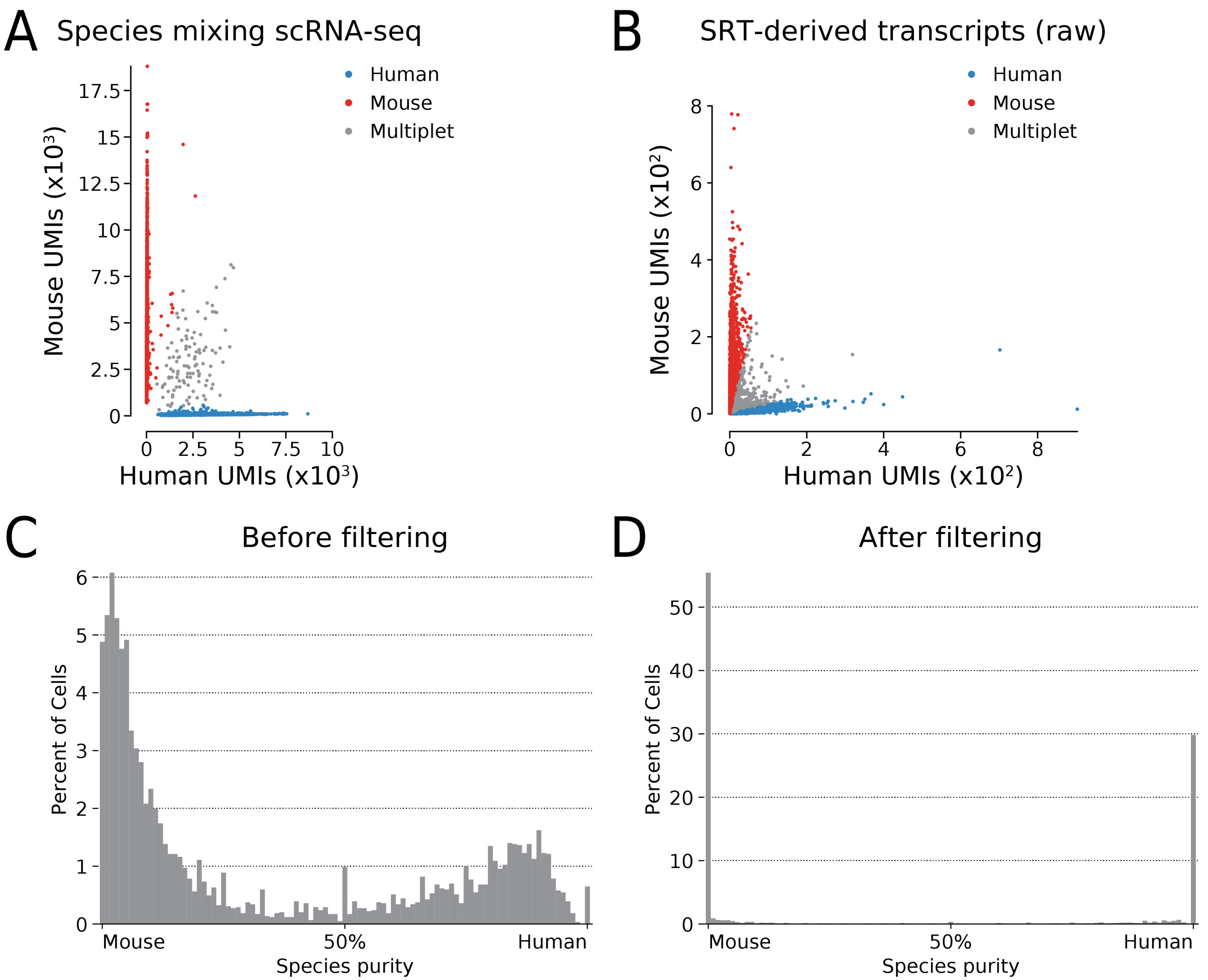


**Supplemental Figure 8: Filtering single cell SRTs reduces intermolecular artifacts.** (A) Barnyard plot from scRNA-seq of HCT-116 and N2a cells shows clean resolution of cell types. Cells were assigned as human or mouse if at least 80% of transcripts in each cell mapped to hg38 or mm10, respectively, or a multiplet otherwise. The multiplet rate was estimated to be 3.2%. (B) Barnyard plot from scCC of HCT-116 and N2a cells without filtering, where an estimated 25.1% of cells were multiplets. (C) Distribution of species purity from unfiltered scCC data. The *x*-axis is the proportion of transcripts mapping to the human or mouse genomes. (D) Distribution of species purity after filtering scCC data. UMI: unique molecular indexes.

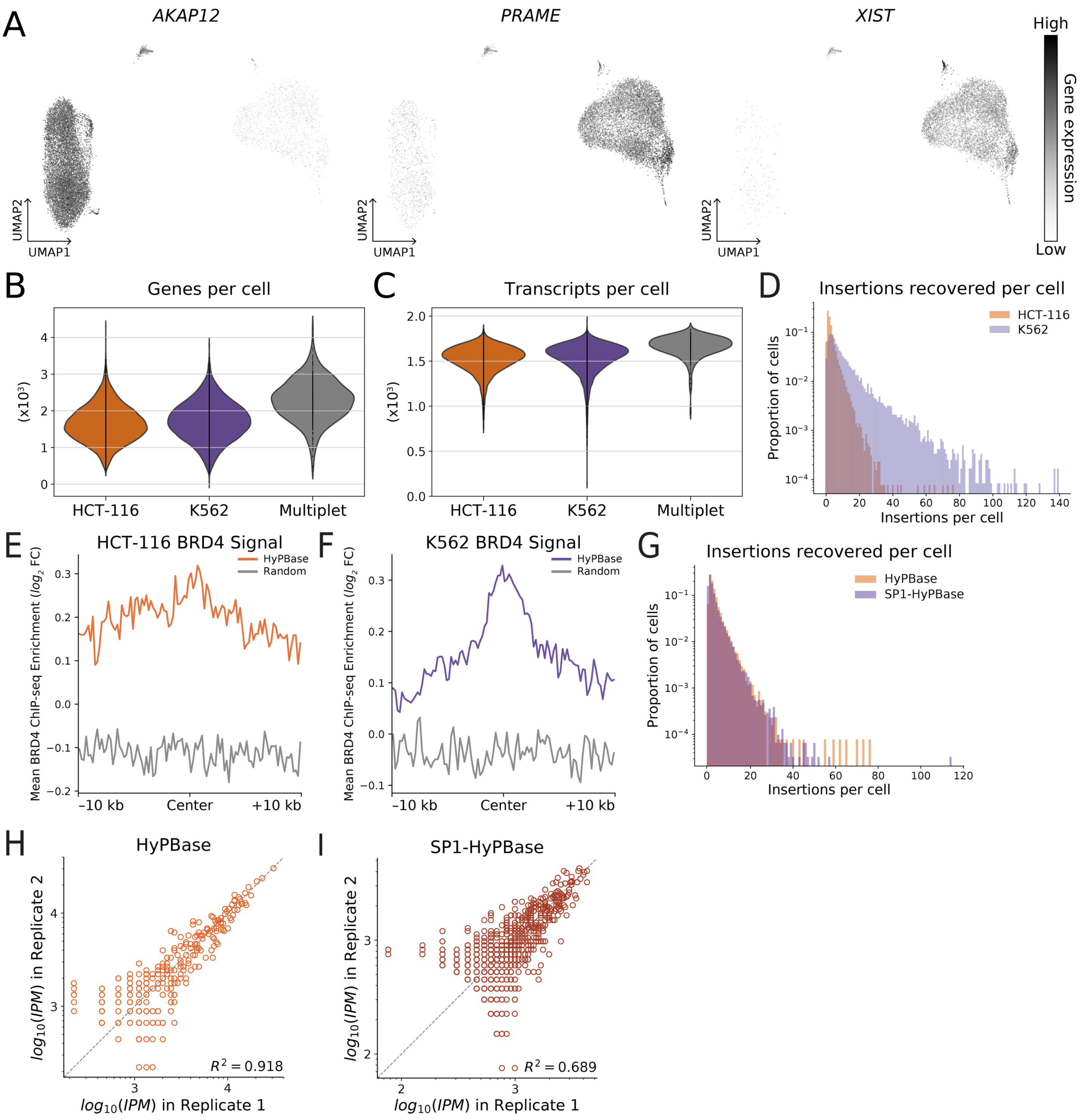


**Supplemental Figure 9: Validation and performance of *in vitro* single cell calling cards.** (A) Expression of three marker genes (*AKAP12, PRAME, XIST)* identify HCT-116 ($n$ = 12,891) and K562 ($n$ = 11,912) cells from scRNA-seq libraries. (B) Distributions of genes per cell by cell type. (C) Distributions of transcripts per cell by cell type. The numbers of genes and transcripts detected per cell were comparable between HCT-116 and K562 cells. (D) Distributions of recovered HyPBase insertions in HCT-116 and K562 cells. (E) Mean BRD4 ChIP-seq signal at HyPBase peaks in HCT-116 cells compared to randomly permuted peaks ($p < 10^{-9}$, KS test). (F) Mean BRD4 ChIP-seq signal at HyPBase peaks in K562 cells compared to randomly permuted peaks ($p < 10^{-9}$, KS test). (G) Distributions of recovered insertions in HCT-116 cells transfected with PB-SRT-Puro and either HyPBase or SP1-HyPBase. (H) Reproducibility of normalized insertions deposited by HyPBase and recovered by scCC at BRD4 binding sites in HCT-116 cells. (I) Reproducibility of normalized insertions deposited by SP1-HyPBase and recovered by scCC at SP1 binding sites in HCT-116 cells. KS: Kolmogorov-Smirnov.

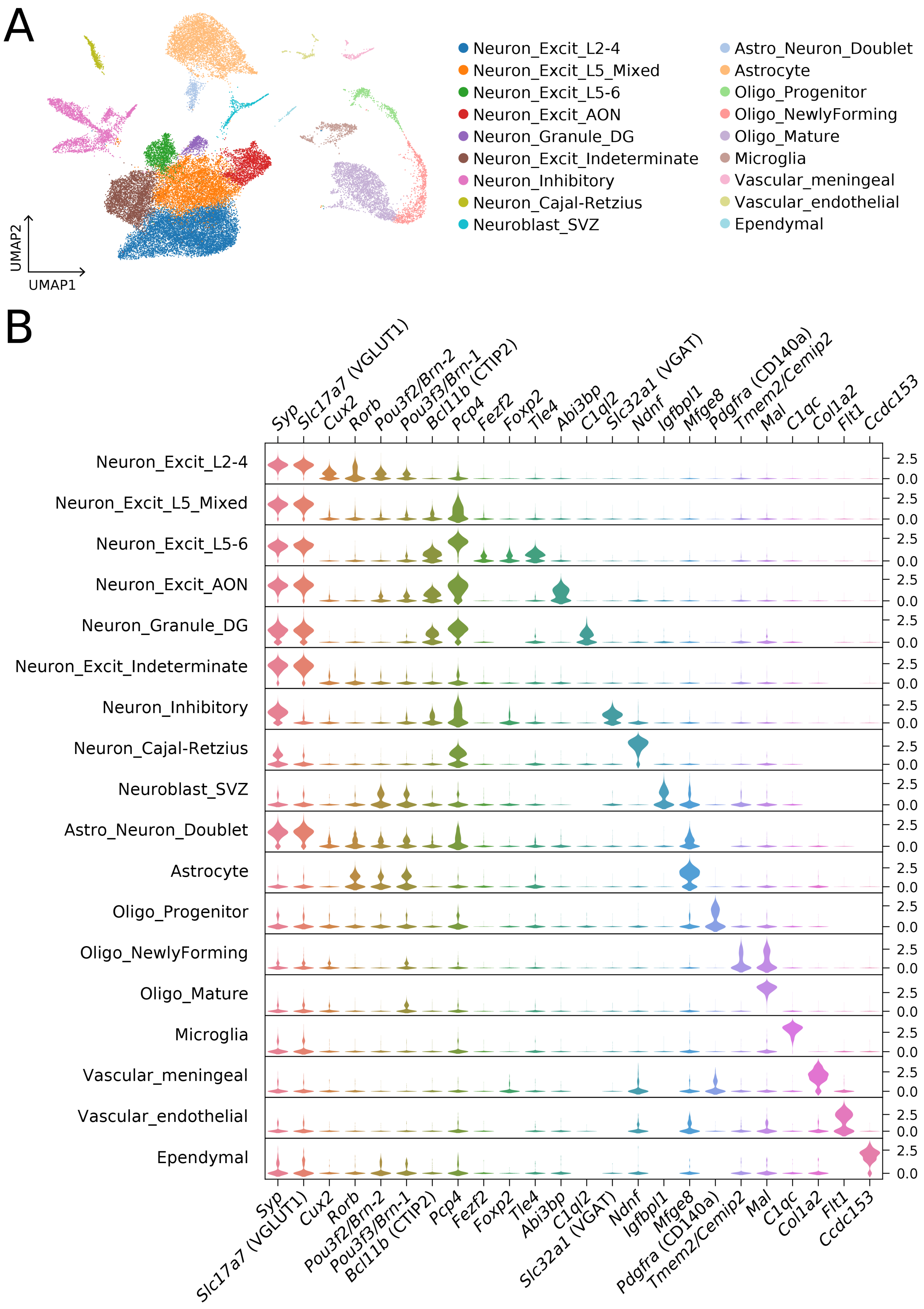

A
Neuron_Excit_L2-4
Neuron_Excit_L5_Mixed
Neuron_Excit_L5-6
Neuron_Excit_AON
Neuron_Granule_DG
Neuron_Excit_Indeterminate
Neuron_Inhibitory
Neuron_Cajal-Retzius
Neuroblast_SVZ
Astro_Neuron_Doublet
Astrocyte
Oligo_Progenitor
Oligo_NewlyForming
Oligo_Mature
Microglia
Vascular_meningeal
Vascular_endothelial
Ependymal
UMAP2
UMAP1
B
Syp
Slc17a7 (VGLUT1)
Cux2
Rorb
Pou3f2/Brn-2
Pou3f3/Brn-1
Bcl11b (CTIP2)
Pcp4
Fezf2
Foxp2
Tle4
Abi3bp
C1ql2
Slc32a1 (VGAT)
Ndnf
Igfbpl1
Mfge8
Pdgfra (CD140a)
Tmem2/Cemip2
Mal
C1qc
Col1a2
Flt1
Ccdc153
2.5
0.0

**Supplemental Figure 10: Clustering of SRT-treated cortical cells and associated marker genes.** (A) Two-dimensional embedding of nine mouse brain (P14-28; 8 cortical, 1 hippocampal) scRNA-seq libraries. Mice were transduced with AAV9-PB-SRT-tdTomato and AAV9-HyPBase at P0-2. Unsupervised dimensionality reduction and Louvain clustering analysis identified 18 populations. (B) Expression profiles of selected marker genes used to identify individual cell types. Excit: excitatory; oligo: oligodendrocytes; SVZ: sub-ventricular zone; AON: anterior olfactory nucleus; DG: dentate gyrus; L2-4: layer 2-4; L5-6: layer 5-6.

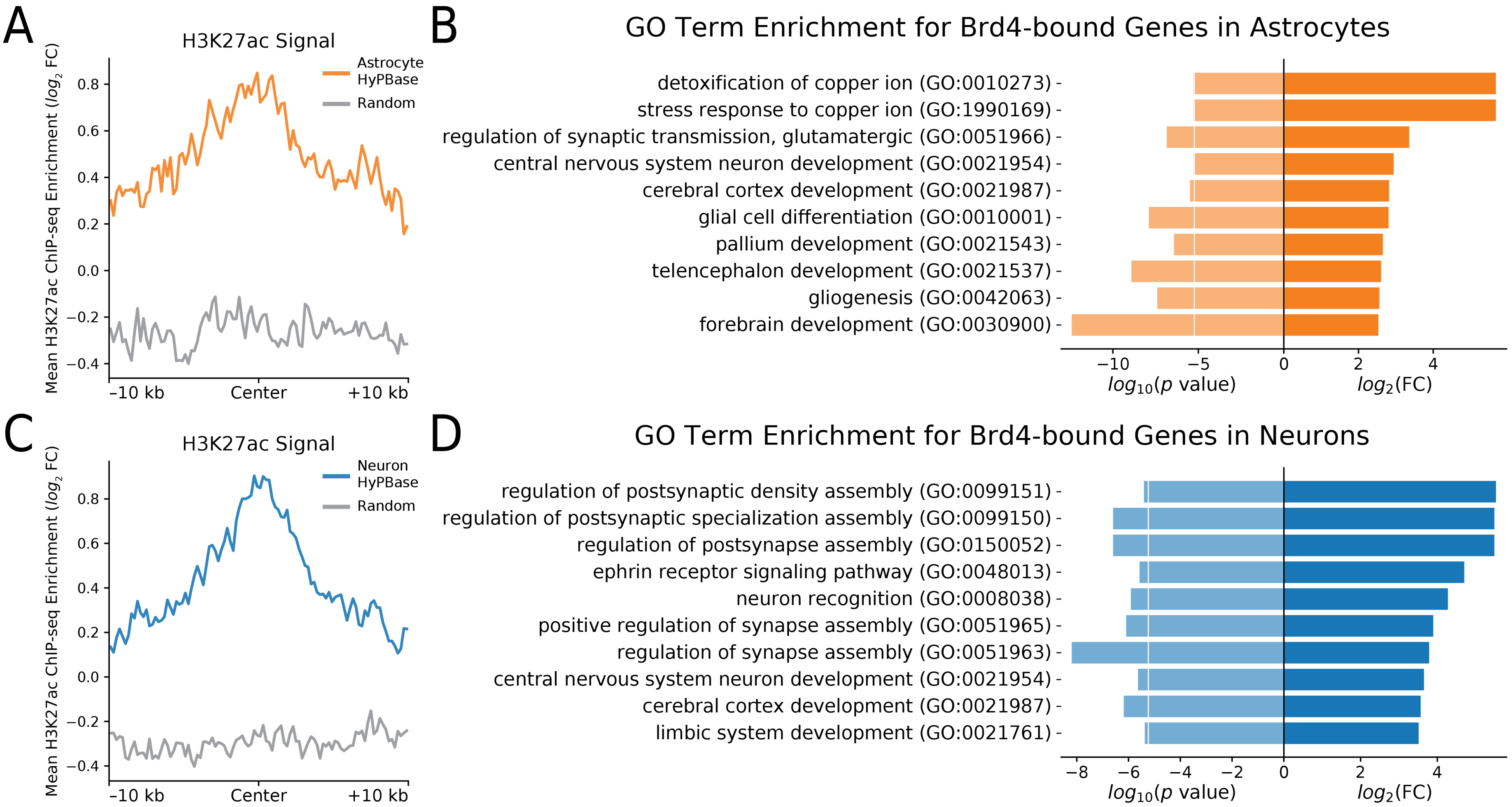


**Supplemental Figure 11: Validation of Brd4 binding *in vivo* in astrocytes and neurons.** (A) Brd4 binding sites, as inferred from astrocytic HyPBase peaks, show significant enrichment for H3K27ac over randomly permuted peaks ($p < 10^{-9}$, KS test). (B) GO term enrichment analysis of genes near astrocytic Brd4 binding sites identifies glial-specific biological processes. The white line indicates the Bonferroni-adjusted *p*-value threshold at $\alpha = 0.05$. (C) Brd4 binding sites from neuronal HyPBase peaks show significant enrichment for H3K27ac over randomly permuted peaks ($p < 10^{-9}$, KS test). (D) GO term enrichment analysis of genes near neuronal Brd4 binding sites identifies neuron-specific biological processes. The white line indicates the Bonferroni-adjusted *p*-value threshold at $\alpha = 0.05$. GO: Gene Ontology; KS: Kolmogorov-Smirnov; FC: fold change.

| Emission | CTCF | H3K9me2 | H3K9me3 | H3K27me3 | H3K36me3 | H4K20me1 | H3K4me1 | H3K4me2 | H3K4me3 | H3K27ac | H3K9ac | H3K79me2 | Candidate state annotation |
|---|---|---|---|---|---|---|---|---|---|---|---|---|---|
| 1 | 86 | 1 | 1 | 1 | 1 | 2 | 13 | 21 | 1 | 5 | 0 | 1 | Insulator |
| 2 | 23 | 1 | 1 | 2 | 0 | 1 | 38 | 97 | 87 | 20 | 29 | 3 | Promoter |
| 3 | 25 | 0 | 0 | 0 | 2 | 1 | 35 | 100 | 100 | 96 | 98 | 37 | |
| 4 | 14 | 0 | 1 | 0 | 2 | 1 | 89 | 98 | 36 | 95 | 41 | 4 | Enhancer |
| 5 | 3 | 2 | 2 | 1 | 1 | 2 | 59 | 57 | 1 | 9 | 1 | 3 | |
| 6 | 3 | 1 | 2 | 0 | 14 | 9 | 77 | 90 | 27 | 53 | 22 | 92 | |
| 7 | 1 | 1 | 4 | 0 | 13 | 10 | 11 | 1 | 0 | 5 | 0 | 85 | Transcribed |
| 8 | 0 | 1 | 3 | 0 | 14 | 5 | 4 | 0 | 0 | 2 | 0 | 12 | |
| 9 | 5 | 0 | 2 | 0 | 32 | 8 | 16 | 5 | 0 | 65 | 1 | 36 | |
| 13 | 0 | 3 | 6 | 1 | 1 | 3 | 4 | 0 | 0 | 0 | 0 | 2 | |
| 12 | 0 | 1 | 19 | 1 | 1 | 3 | 3 | 0 | 1 | 0 | 0 | 1 | Repressed |
| 15 | 0 | 2 | 3 | 30 | 0 | 5 | 3 | 0 | 0 | 0 | 0 | 1 | |
| 11 | 0 | 0 | 0 | 0 | 0 | 0 | 0 | 0 | 0 | 0 | 0 | 0 | Inactive |
| 14 | 0 | 1 | 1 | 1 | 0 | 1 | 2 | 0 | 0 | 0 | 0 | 0 | |
| 10 | 3 | 48 | 69 | 30 | 54 | 36 | 28 | 21 | 34 | 21 | 15 | 28 | |

Chromatin states (rows); Chromatin mark observation frequency (%)

**Supplemental Table 1: ChromHMM's chromatin state annotations in HCT-116 cells.**

| Cluster | # cells | # insertions | Mean |
|---|---|---|---|
| Astrocyte | 4,727 | 17,102 | 3.6 |
| Astro_Neuron_Doublet | 394 | 1,667 | 4.2 |
| Ependymal | 107 | 153 | 1.4 |
| Microglia | 569 | 246 | 0.4 |
| Neuroblast_SVZ | 369 | 1,095 | 3.0 |
| Neuron_Cajal-Retzius | 552 | 4,415 | 8.0 |
| Neuron_Excit_AON | 1,939 | 8,328 | 4.3 |
| Neuron_Excit_Indeterminate | 3,660 | 6,474 | 1.8 |
| Neuron_Excit_L2-4 | 9,083 | 30,225 | 3.3 |
| Neuron_Excit_L5 | 5,544 | 27,117 | 4.9 |
| Neuron_Excit_L6 | 1,436 | 5,317 | 3.7 |
| Neuron_Granule_DG | 535 | 1,733 | 3.2 |
| Neuron_Inhibitory | 2,409 | 6,690 | 2.8 |
| Oligo_Mature | 2,740 | 1,781 | 0.7 |
| Oligo_NewlyForming | 959 | 680 | 0.7 |
| Oligo_Progenitor | 504 | 488 | 1.0 |
| Vascular_endothelial | 196 | 71 | 0.4 |
| Vascular_meningeal | 227 | 277 | 1.2 |
| **Total** | **35,950** | **113,859** | **3.2** |

**Supplemental Table 2: Breakdown of cortical cell types and scCC HyPBase insertions per cluster.** SVZ: sub-ventricular zone; AON: anterior olfactory nucleus; DG: dentate gyrus; L2-4: layer 2-4; L5-6: layer 5-6.

| Plasmid | Description | Internal Accession No. | Addgene? | Addgene Accession No. |
|---|---|---|---|---|
| PBase | *piggyBac* transposase | pRM1024 | NA | NA |
| HyPBase | Hyperactive *piggyBac* transposase | pRM1114 | NA | NA |
| SP1-PBase | SP1 fused to *piggyBac* | pRM1023 | NA | NA |
| SP1-HyPBase | SP1 fused to hyperactive *piggyBac* | pRM1677 | NA | NA |
| PB-SRT-Puro | *piggyBac* SRT with puromycin reporter | pRM1304 | NA | NA |
| PB-SRT-tdTomato | *piggyBac* SRT with tdTomato reporter | pRM1535 | NA | NA |
| SB100X | Hyperactive *Sleeping Beauty* transposase | pRM1137 | NA | NA |
| SB-SRT-Puro | *Sleeping Beauty* SRT with puromycin reporter | pRM1668 | NA | NA |
| AAV-HyPBase | Hyperactive *piggyBac* transposase in an AAV vector | pRM1217 | NA | NA |
| AAV-PB-SRT-tdTomato | PB-SRT-tdTomato in an AAV vector | pRM1648 | NA | NA |

**Supplemental Table 3: Plasmids referenced in this work**

| Primer | Primer Sequence | Purification | Notes |
|---|---|---|---|
| SMART_dT18VN | AAGCAGTGGTATCAACGCAGAGTACGTTTTTTTTTTTTTTTTTTTTTTTTTTTTTTTVN | Standard desalt | RT primer for bulk RNA calling card recovery |
| SMART | AAGCAGTGGTATCAACGCAGAGT | Standard desalt | PCR primer for bulk RNA calling card amplification |
| SRT_PAC_F1 | CAACCTCCCCTTCTACGAGC | Standard desalt | Puromycin marker in SRT |
| SRT_tdTomato_F1 | TCCTGTACGGCATGGACGAG | Standard desalt | tdTomato marker in SRT |
| Raff_ACTB_F | CCTCGCCTTTGCCGATCCG | Standard desalt | Human ACTB primer (for RT control) |
| Raff_ACTB_R | GGATCTTCATGAGGTAGTCAGTCAGGTCC | Standard desalt | Human ACTB primer (for RT control) |
| OM-PB-ACG | AATGATACGGCGACCACCGAGATCTACACTCTTTCCCTACACGACGCTCTTCCGATCTACGTTTACGCAGACTATCTTTCTAG | Standard desalt | For use with *piggyBac* SRTs |
| OM-PB-CTA | AATGATACGGCGACCACCGAGATCTACACTCTTTCCCTACACGACGCTCTTCCGATCTCTATTTACGCAGACTATCTTTCTAG | Standard desalt | For use with *piggyBac* SRTs |
| OM-PB-GAT | AATGATACGGCGACCACCGAGATCTACACTCTTTCCCTACACGACGCTCTTCCGATCTGATTTTACGCAGACTATCTTTCTAG | Standard desalt | For use with *piggyBac* SRTs |
| OM-PB-TGC | AATGATACGGCGACCACCGAGATCTACACTCTTTCCCTACACGACGCTCTTCCGATCTTGCTTTACGCAGACTATCTTTCTAG | Standard desalt | For use with *piggyBac* SRTs |
| OM-PB-TAG | AATGATACGGCGACCACCGAGATCTACACTCTTTCCCTACACGACGCTCTTCCGATCTTAGTTTACGCAGACTATCTTTCTAG | Standard desalt | For use with *piggyBac* SRTs |
| OM-PB-ATC | AATGATACGGCGACCACCGAGATCTACACTCTTTCCCTACACGACGCTCTTCCGATCTATCTTTACGCAGACTATCTTTCTAG | Standard desalt | For use with *piggyBac* SRTs |
| OM-PB-CGT | AATGATACGGCGACCACCGAGATCTACACTCTTTCCCTACACGACGCTCTTCCGATCTCGTTTTACGCAGACTATCTTTCTAG | Standard desalt | For use with *piggyBac* SRTs |
| OM-PB-GCA | AATGATACGGCGACCACCGAGATCTACACTCTTTCCCTACACGACGCTCTTCCGATCTGCATTTACGCAGACTATCTTTCTAG | Standard desalt | For use with *piggyBac* SRTs |
| OM-SB-ACG | AATGATACGGCGACCACCGAACACTCTTTCCCTACACGACGCTCTTCCGATCTACGTAAGTGTATGTAAACTTCCGACTTCAA | Standard desalt | For use with *Sleeping Beauty* SRTs |
| OM-SB-CTA | AATGATACGGCGACCACCGAACACTCTTTCCCTACACGACGCTCTTCCGATCTCTATAAGTGTATGTAAACTTCCGACTTCAA | Standard desalt | For use with *Sleeping Beauty* SRTs |
| OM-SB-GAT | AATGATACGGCGACCACCGAACACTCTTTCCCTACACGACGCTCTTCCGATCTGATTAAGTGTATGTAAACTTCCGACTTCAA | Standard desalt | For use with *Sleeping Beauty* SRTs |
| OM-SB-TGC | AATGATACGGCGACCACCGAACACTCTTTCCCTACACGACGCTCTTCCGATCTTGCTAAGTGTATGTAAACTTCCGACTTCAA | Standard desalt | For use with *Sleeping Beauty* SRTs |
| OM-SB-TAG | AATGATACGGCGACCACCGAACACTCTTTCCCTACACGACGCTCTTCCGATCTTAGTAAGTGTATGTAAACTTCCGACTTCAA | Standard desalt | For use with *Sleeping Beauty* SRTs |
| OM-SB-ATC | AATGATACGGCGACCACCGAACACTCTTTCCCTACACGACGCTCTTCCGATCTATCTAAGTGTATGTAAACTTCCGACTTCAA | Standard desalt | For use with *Sleeping Beauty* SRTs |

| | | | |
|---|---|---|---|
| OM-SB-CGT | AATGATACGGCGACCACCGAACACTCTTTCCCTACACGACGCTCTTCCGATCTCGTTAAGTGTATGTAAACTTCCGACTTCAA | Standard desalt | For use with *Sleeping Beauty* SRTs |
| OM-SB-GCA | AATGATACGGCGACCACCGAACACTCTTTCCCTACACGACGCTCTTCCGATCTGCATAAGTGTATGTAAACTTCCGACTTCAA | Standard desalt | For use with *Sleeping Beauty* SRTs |
| N7 indexed primer | CAAGCAGAAGACGGCATACGAGAT[index]GTCTCGTGGGCTCGG | Standard desalt | Uniquely identifies each bulk RNA calling card library in conjunction with barcoded transposon primer |
| 10x_TSO | AAGCAGTGGTATCAACGCAGAGTACATrGrGrG | Standard desalt | For continuing 10x scRNA-seq prep after splitting first RT product in half |
| Bio_Illumina_Seq1_scCC_10X_3xPT | /5Phos/ACACTCTTTCCC/iBiodT/ACACGACGCTCTTCCGA*T*C*T | HPLC | Single cell calling card primer for use with 10x Chromium 3' v2 kit |
| Bio_Long_PB_LTR_3xPT | /5Phos/GCGTCAATTTTACGCAGAC/iBiodT/ATCTTTC*T*A*G | HPLC | Single cell calling card primer for use with *piggyBac* SRTs |
| scCC_P5_adapter | AATGATACGGCGACCACCGAGATCTTCACTCATTCCACACGACTCCTTGCCAGTCTC*T | Standard desalt | Adapter for scCC (needs to be pre-annealed with scCC_P7_adapter) |
| scCC_P7_adapter | /5Phos/GAGACTGGCAAGTACACGTCGCACTCACCATGA[index]ATCTCGTATGCCGTCTTCTGCTTG | Standard desalt | Adapter for scCC (needs to be pre-annealed with scCC_P5_adapter) |
| scCC_P5_primer | AATGATACGGCGACCACCGAGATC | Standard desalt | For final scCC library PCR |
| scCC_P7_primer | CAAGCAGAAGACGGCATACGAGAT | Standard desalt | For final scCC library PCR |
| scCC_PB_CustomRead2 | CGTGTAGGGAAAGAGTGTGCGTCAATTTTACGCAGACTATCTTTCTAG | PAGE | For custom sequencing of *piggyBac* scCC libraries; read 2 should begin with GGTTAA |
| scCC_CustomIndex1 | GAGACTGGCAAGTACACGTCGCACTCACCATGA | PAGE | For custom sequencing of scCC libraries |

**Supplemental Table 4: Oligonucleotides referenced in this work**

| Target | Cell type | Source | PI | Accession(s) | Control File(s) | DOI |
|---|---|---|---|---|---|---|
| SP1 | HCT-116 | ENCODE | Myers, Richard | ENCFF000PCT | ENCFF000PBO | |
| BRD4 | HCT-116 | Publication | Firestein, Ron | SRR2481799 | SRR2481800 | 10.1172/JCI83265 |
| H3K27ac | HCT-116 | ENCODE | Bernstein, Bradley | ENCFF082JPN, ENCFF176BXC | ENCFF048ZOQ, ENCFF827YXC | |
| H3K4me1 | HCT-116 | ENCODE | Bernstein, Bradley | ENCFF088BWP, ENCFF804MJI | ENCFF048ZOQ, ENCFF827YXC | |
| H3K9me3 | HCT-116 | ENCODE | Bernstein, Bradley | ENCFF578MDZ, ENCFF033XOG | ENCFF048ZOQ, ENCFF827YXC | |
| H3K27me3 | HCT-116 | ENCODE | Bernstein, Bradley | ENCFF281SBT, ENCFF124GII | ENCFF048ZOQ, ENCFF827YXC | |
| CTCF | HCT-116 | ENCODE | Myers, Richard | ENCFF000OZC | ENCFF000PBO | |
| H3K9me2 | HCT-116 | ENCODE | Bernstein, Bradley | ENCFF760OZN, ENCFF565FDP | ENCFF048ZOQ, ENCFF827YXC | |
| H3K36me3 | HCT-116 | ENCODE | Bernstein, Bradley | ENCFF850EAH, ENCFF312RKB | ENCFF048ZOQ, ENCFF827YXC | |
| H4K20me1 | HCT-116 | ENCODE | Bernstein, Bradley | ENCFF070JDY, ENCFF334HHB | ENCFF048ZOQ, ENCFF827YXC | |
| H3K4me2 | HCT-116 | ENCODE | Bernstein, Bradley | ENCFF936MMN, ENCFF937OOL | ENCFF048ZOQ, ENCFF827YXC | |
| H3K4me3 | HCT-116 | ENCODE | Bernstein, Bradley | ENCFF183OZI, ENCFF659FPR | ENCFF048ZOQ, ENCFF827YXC | |
| H3K9ac | HCT-116 | ENCODE | Bernstein, Bradley | ENCFF408RRT | ENCFF413RQG | |
| H3K79me2 | HCT-116 | ENCODE | Bernstein, Bradley | ENCFF865KPW, ENCFF947YPU | ENCFF048ZOQ, ENCFF827YXC | |
| BRD4 | K562 | ENCODE | Bernstein, Bradley | ENCFF335PHG | ENCFF000BWK | |
| H3K27ac | K562 | ENCODE | Bernstein, Bradley | ENCFF000BXH | ENCFF000BWK | |
| H3K27ac | Mouse cortex | Publication | Greenberg, Michael | SRR6129714 | SRR6129695 | 10.1016/j.cell.2017.09.047 |

**Supplemental Table 5: External ChIP-seq datasets referenced in this work**
